# MGAT5 (GnTase-V) products are overexpressed in primary osteoblastic osteogenic sarcoma

**DOI:** 10.1101/2023.04.11.536427

**Authors:** John McClure, Sheena F McClure

**Affiliations:** Professor Emeritus of Pathology, University of Manchester Medical School, UK; Research Associate (Rtd), University of Manchester Medical School, UK

**Keywords:** MGAT5 (GnTase-V), N-glycosylation, 0-glycosylation, Primary Osteoblastic Osteogenic Sarcoma, Osteoclasts, Endothelial Cells, Mast Cells, Neoangiogenesis

## Abstract

It is well documented that MGAT5 (Glycosyltransferase V) products are over expressed in carcinomas and likely accelerate metastasis by promoting neoangiogenesis. We now report a similar over expression in primary osteoblastic osteogenic sarcoma and demonstrate new intratumoural blood vessels. Expansion of the tumours is by osteoclastic resorption of both trabecular and cortical bone with the latter by cutting channels similar to physiological cutting cones. Osteoclasts, adjacent capillaries, and related mast cells also overexpress MGAT5 products and we speculate that this facilitates neoangiogenesis.

## INTRODUCTION

It is well documented that N-acetylglucosaminyltransferase-V (GnTase-V or MGAT5) alters the structure of specific N-glycans by modifying α1-6 linked mannose with a β1-6 linked N-acetylglucosamine branch. It is also well documented that β1-6 branching is over expressed on cancer cell surfaces and is believed to accelerate metastasis by promoting angiogenesis. We have recently reported β1-6 branching over-expression in chondrosarcoma seemingly the first such report (1). Similar changes do not appear to have been described for other sarcomas. We have recently had the opportunity of studying a number of osteoblastic osteogenic osteosarcomas and our findings form the basis of this report.

## MATERIALS AND METHODS

Six examples of osteoblastic osteogenic sarcoma were available in a pathology archive – five located at the lower end of femur and one at the upper end of tibia. All were from individuals in the second and third decades of life. The resection specimens had been photographed and coronal one centimetre thick sections made on a bandsaw. Contact radiographs were prepared from these. Multiple tissue blocks had been decalcified under x-ray control, embedded in paraffin wax and sections cut for histological examination. Staining was haematoxylin and eosin (H&E) and by lectin histochemistry for *Phaseolus vulgaris* lectin (l-PHA) and *Psophocarpus tetragonolobus* lectin (PTL-II).

## RESULTS

The photomacrographs and the contact radiographs showed tumours eccentrically placed near the ends of long bones. Intratumoural calcification was variable. Generally, the margins were cleared with dissolution of the trabecular bone. The eccentric position meant that tumour impinged on adjacent cortical bone with varying degree of disruption of the latter.

Histologically, there was new “malignant” osteoid formation by tumour cells. This osteoid material showed varying degrees of mineralization. Resorption of native trabecular bone was by osteoclasts separated from tumour by a connective tissue. Resorption of cortical bone was by cutting channels reminiscent of osteonal cutting cones. Indeed, there was expansion of some Haversian canals suggestive of a beginning cutting cone/channel. These latter were inevitably led by osteoclasts with trailing capillaries and mesenchymal tissue containing mast cells.

Osteosarcoma cells expressed high levels of l-PHA ligands. This expression was cytoplasmic and membranous in location and never intranuclear. Tumour tissue was highly vascular as demonstrated by the expression of PTL-II ligands. These blood vessels showed variation in shape and size. At the tumour margins mesenchymal tissue was infiltrated by mast cells which were never seen within tumour tissue. Mast cells also expressed ligands for l-PHA.

Osteoclasts located both on trabecular bone and in cutting channels expressed l-PHA ligands. These were cytoplasmic and membranous (never intranuclear) in location. The expression was particularly marked at the resorbing surface. Within the cutting channels mast cells and endothelial cells expressed l-PHA ligands. Only endothelial cells expressed PTL-II ligands.

## DISCUSSION

The Human Protein Atlas (https://www.proteinatlas.org) on its cancer tissue page lists some twenty human malignancies (mostly carcinomas and also including malignant melanoma and lymphoma) in which there is overexpression of MGAT5. Sarcoma is not in the list. We have previously reported such overexpression in chondrosarcoma (1) and now, herein, osteosarcoma.

We have applied lectin histochemistry using the lectins l-PHA (*Phaseolus vulgaris*) and PTL-II (*Phosocarpus tetragonolobus*) in the present study. The ligand for l-PHA is the beta 1-6 linkage in complex N-glycans which is initiated by the Golgi-bound glycosyltransferase GnTaseV/MGAT5. The demonstration of these linkages in lectin histochemical reactions is an index of MGAT5 activity.

Amounts of MGAT5 glycan products are commonly increased in malignancies and appear to be correlated with disease progression by promoting both local tumour proliferation and metastatic spread probably by promoting angiogenesis (2). It has been postulated that MGAT5 selectively remodels the endothelial cell surface by the regulated binding of Galectin-1 (Gal-1) which on recognition of complex N-glycans on VEGFR2 activates VEGF signalling (3,4).

The significance of MGAT5 overexpression in malignancy is in the potential for developing therapeutic targets. It has been shown that the MGAT5 – acceptor complex structure suggests a catalytic mechanism, explaining the known inhibition of MGAT5 by GnT-III another branching enzyme and this provides a basis for the rational design of drugs targeting N-glycans branching (5).

The osteosarcomas in this study had a high degree of intratumoural vascularity. This is in contrast to chondrosarcoma where blood vessels are distributed around the periphery of tumour nodules. The intraosteosarcoma blood vessels universally stained with the lectin PTL-II. This reaction is an index of O-glycosylation which is apparently necessary for the development and maintenance of the integrity of blood vessels (6,7). The biosynthesis of this core 1 O-glycan (Galβ 1-3 GalNAcα1-Ser/Thr, T antigen) is controlled by core 1 β1-3-galactosyltransferase (T-synthase) catalysing the addition of Gal to GalNAcα1-Ser/Thr (Tn antigen) which is the reaction recognised by PTL-II. Tn antigen is a precursor for extended and branched O-glycans. Disruption of these processes again offers therapeutic potential since the formation of competent intratumoural blood vessels is clearly necessary for metastatic spread.

It is notable that the osteosarcomas were eccentrically located within bone and, therefore, closer to an adjacent cortical surface. Tumour expansion had occurred by osteoclastic resorption of both trabecular and cortical bone tissue. Osteoclasts were accompanied by a connective tissue containing capillaries and mast cells. Resorption of cortical bone was by cutting channels reminiscent of physiological cutting cones. These channels were led by osteoclasts again with companion capillaries and mast cells. Neoangiogenesis is related to osteoclasis to meet metabolic and catabolic requirements. Osteoclasts, endothelial cells and mast cells were l-PHA positive and we speculate that this facilitates neoangiogenesis by the mechanisms described above. The eccentric location of the tumour renders the proximate cortex vulnerable to osteoclastic attack causing mechanical weakening and susceptibility to pathological fracture.

In osteoclasts the reactivity to l-PHA was present on cell membranes and within cytoplasm but mainly on the former. In many cells the reactivity showed polarity in the resorption zone. This may be the result of membrane folding in the ruffled border. Intriguingly, there is a report that MGAT5 glycan products stimulated membrane ruffling and phosphatidylinositol 3 kinase-protein kinase B activation in mammary carcinoma cells (8). This raises the possibility that MGAT5 N-glycosylation has a role in the creation and/or maintenance of the ruffled border of osteoclasts.

**Figure 1.**
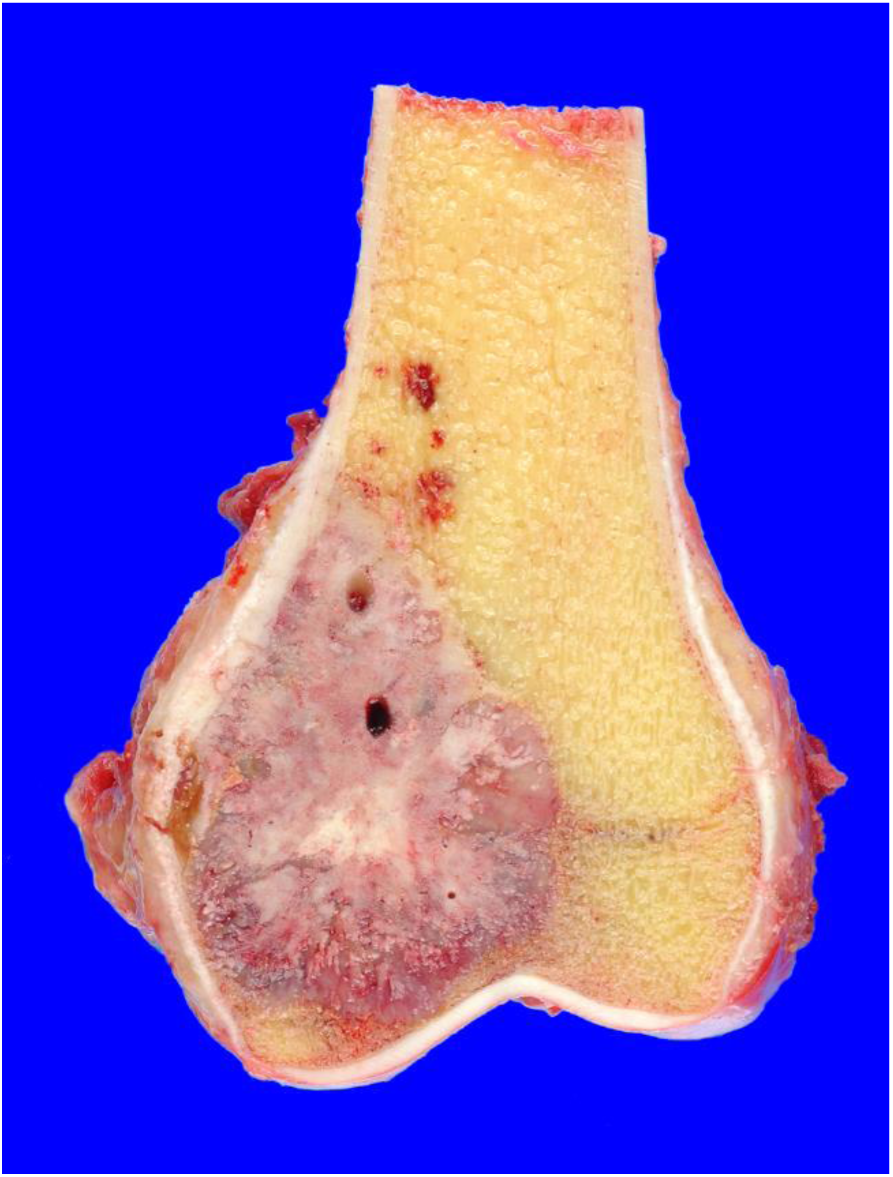
Osteogenic sarcoma in the lower end of femur. Note its eccentric position and disruption of adjacent cortex.

**Figure 2.**
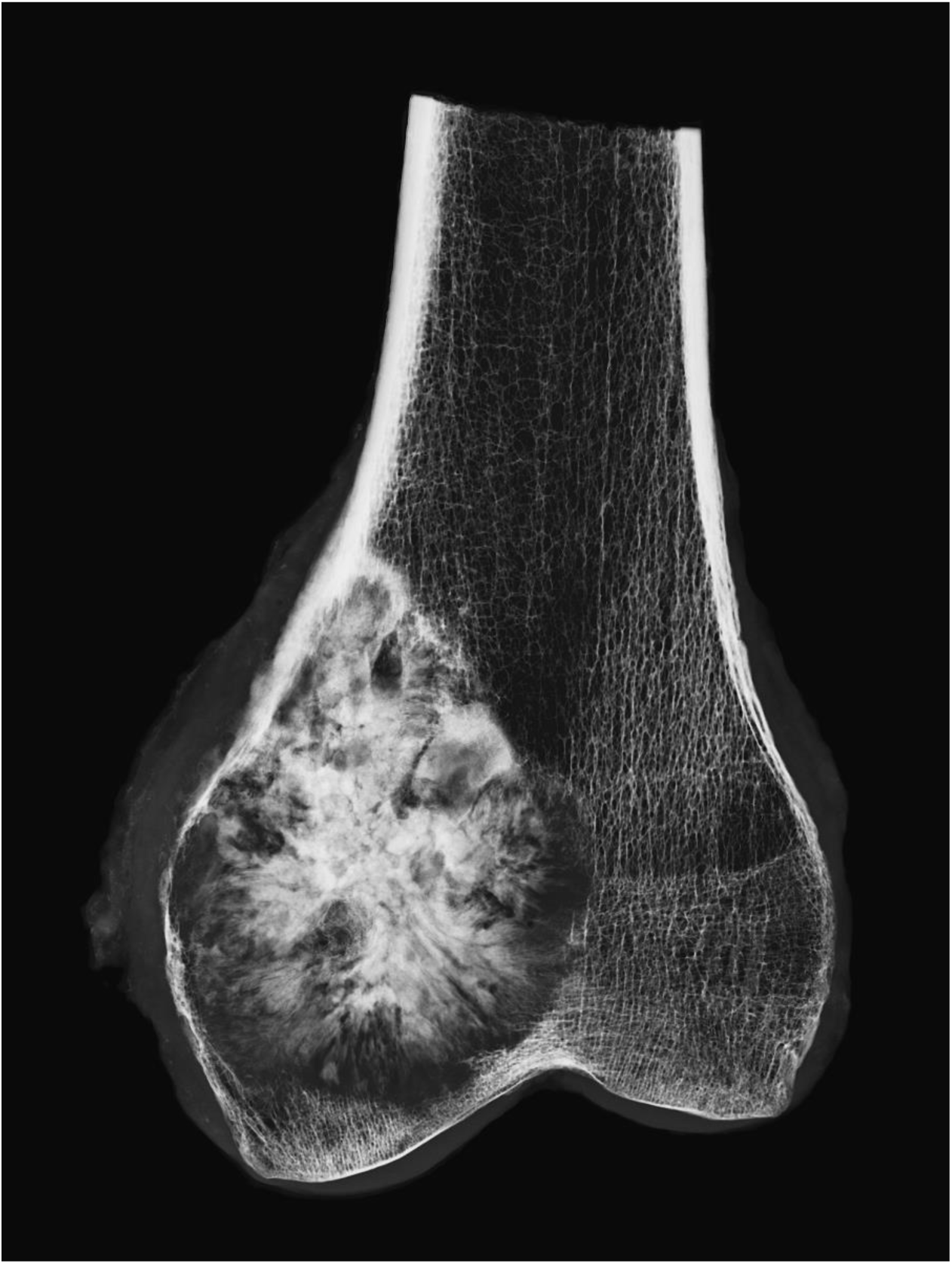
Contact radiograph of sample in Figure 1. Centrally there is calcification of tumour with clear margins. The cortex is thinned and disrupted.

**Figure 3.**
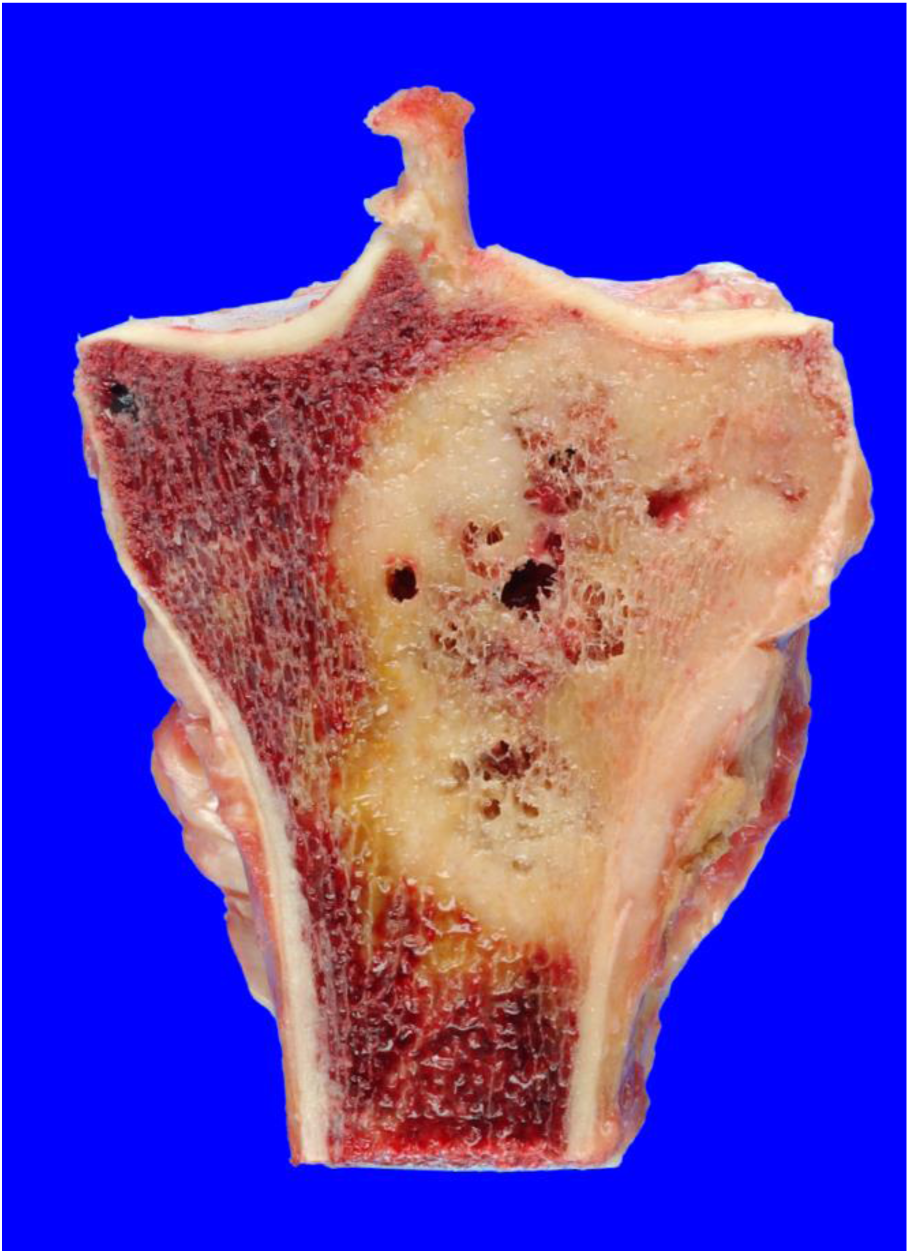
Osteogenic sarcoma in the upper end of tibia. Again it is eccentrically positioned and impinges on the cortex.

**Figure 4.**
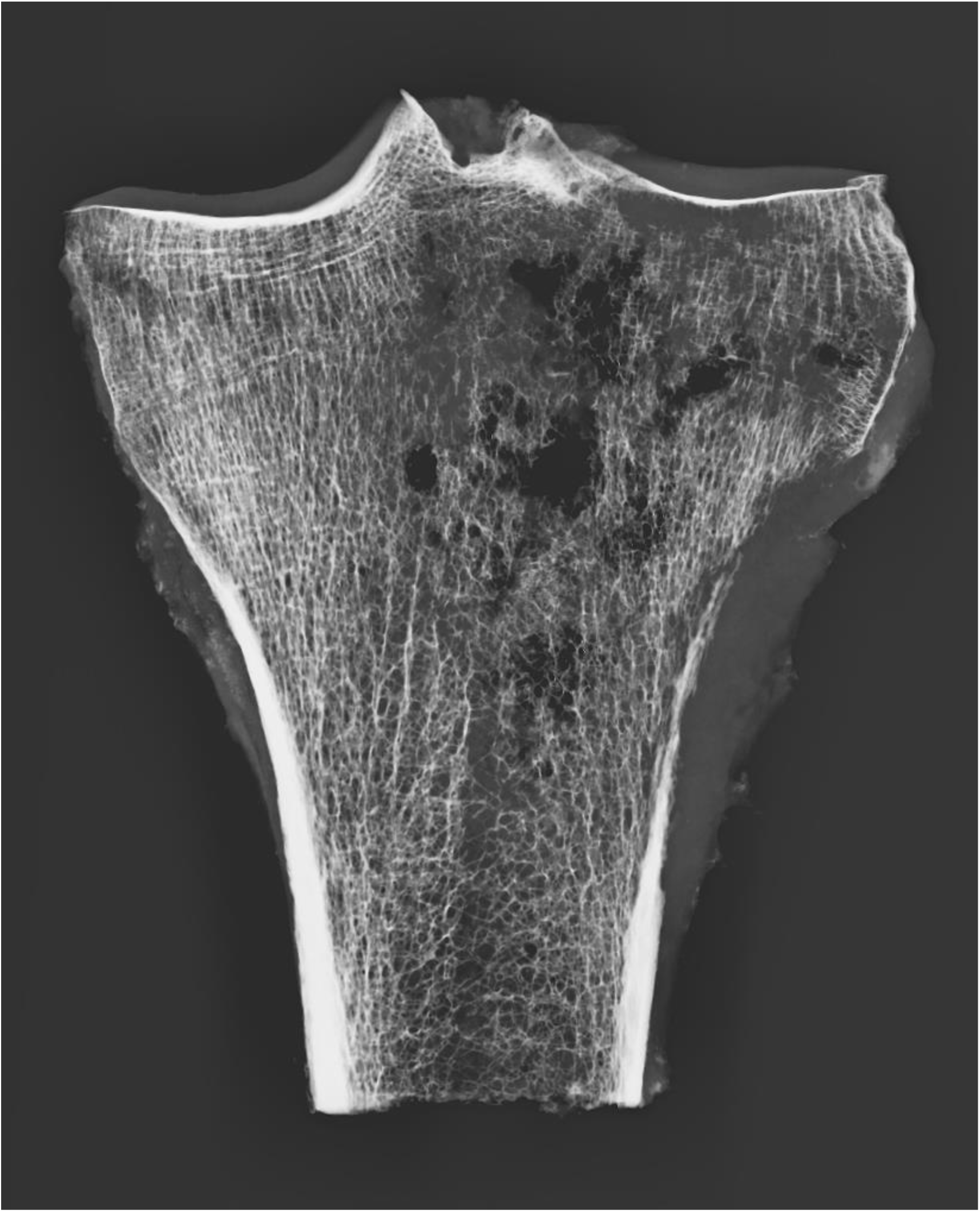
Contact radiograph of sample in Figure 3. The cortex is disrupted.

**Figure 5.**
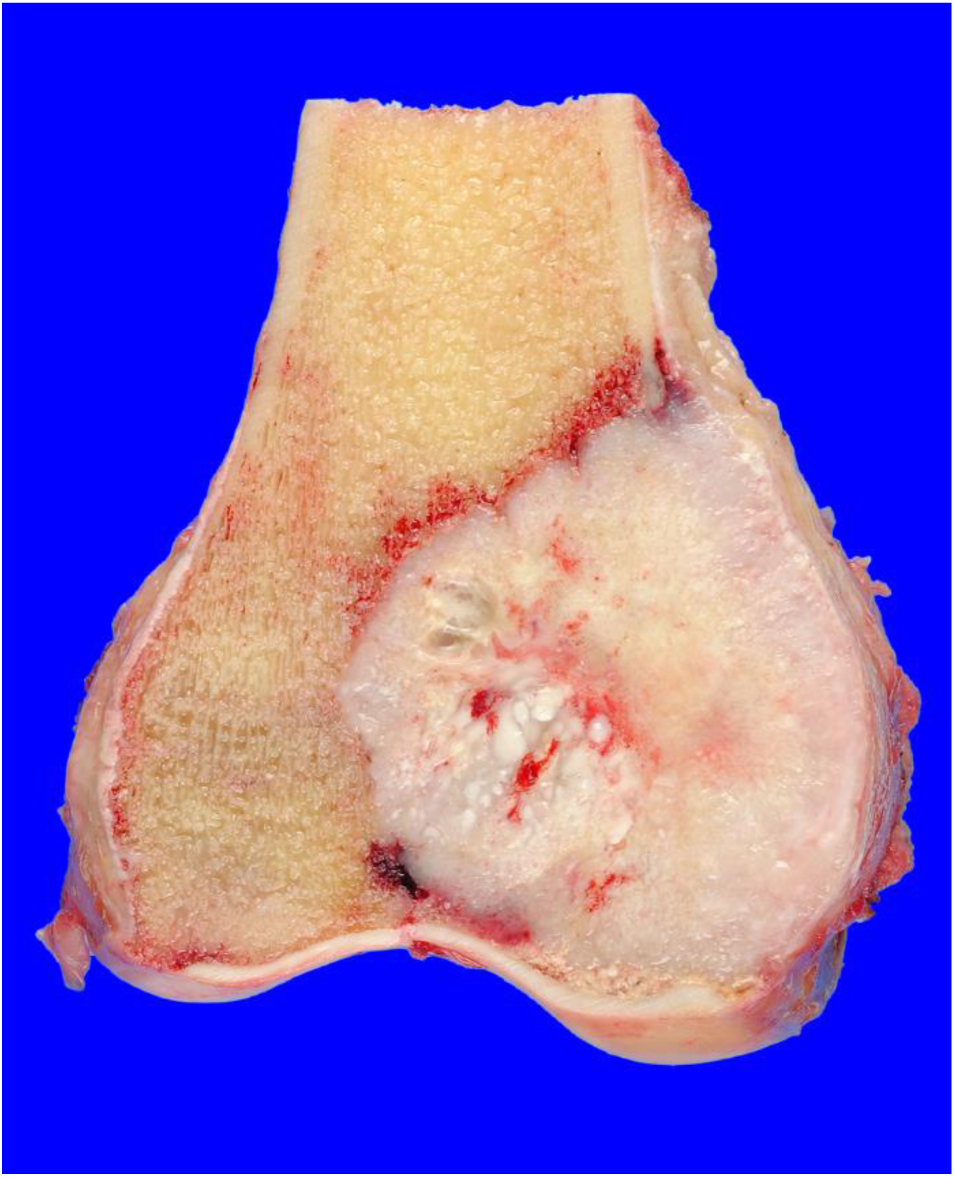
Another osteogenic sarcoma at the lower end of femur. The cortex has been disrupted by tumour.

**Figure 6.**
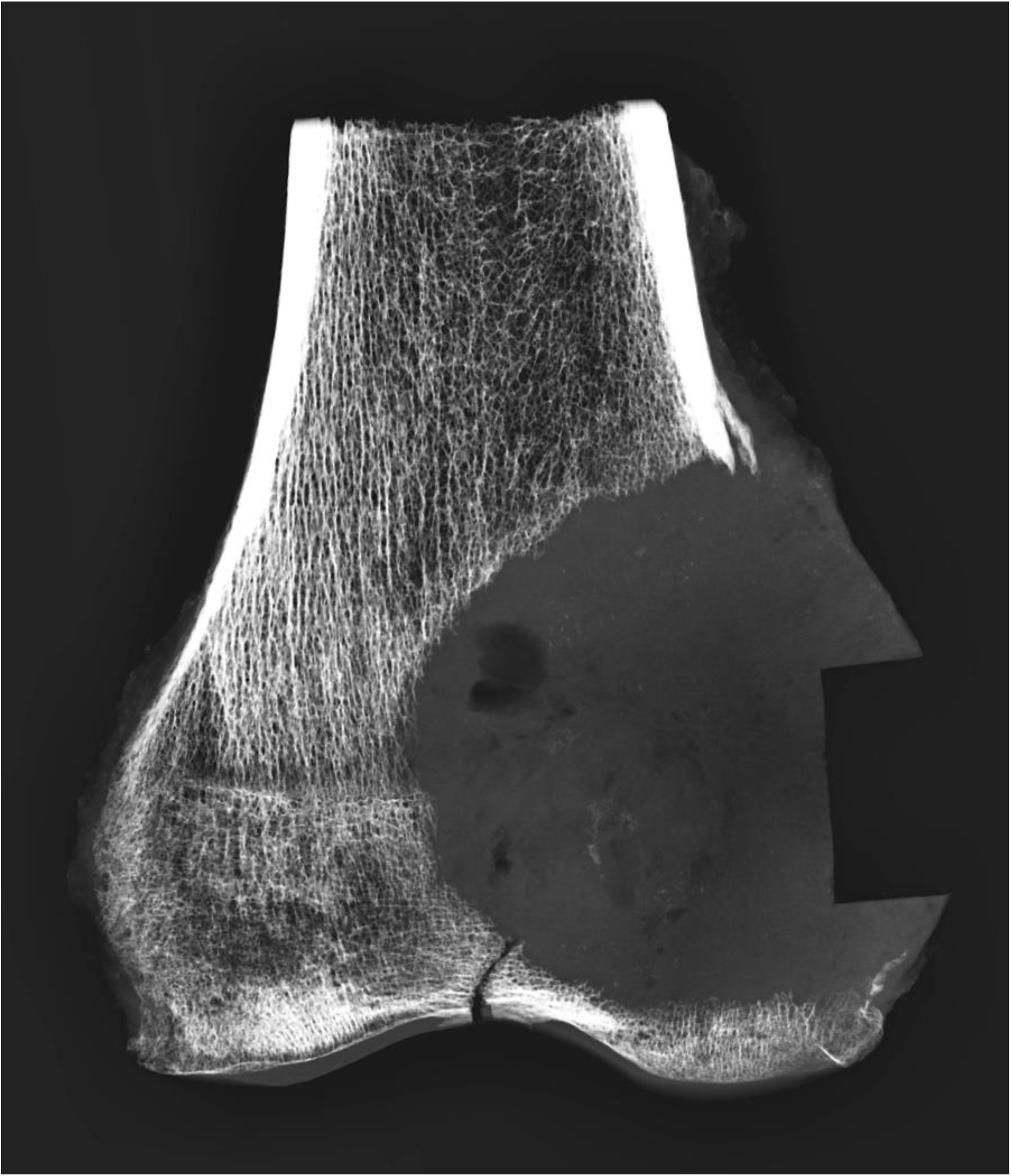
Contact radiograph of sample in Figure 5. The cortex has been completely disrupted by tumour. There is a small amount of calcification near the centre of the lesion. A tissue sample has been taken from the right-hand external margin of the tumour.

**Figure 7.**
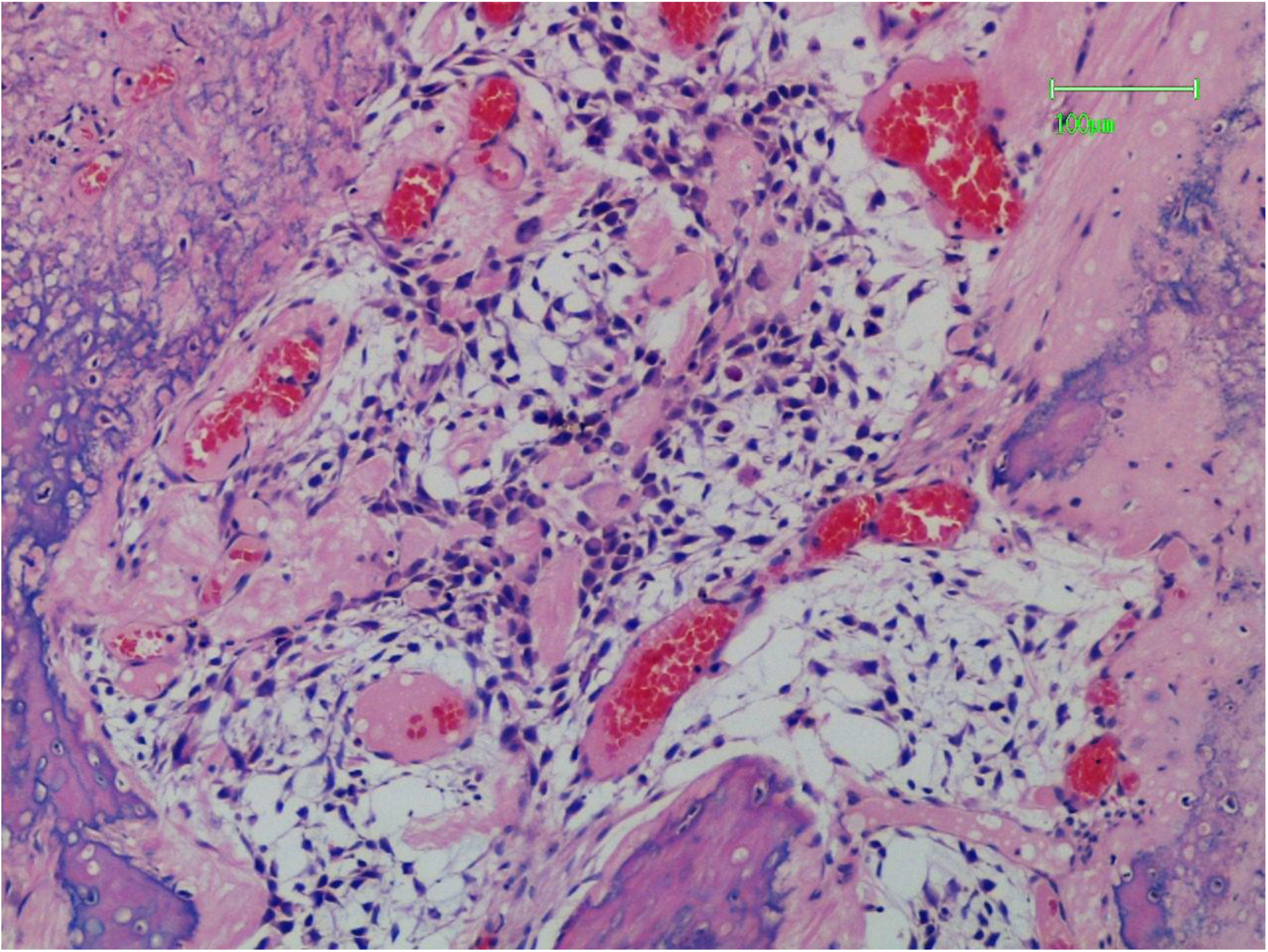
Osteoblastic osteogenic osteosarcoma with central formation of unmineralized tumour osteoid. Peripherally the tumour matrix is calcified (Haematoxylin and Eosin – HE).

**Figure 8.**
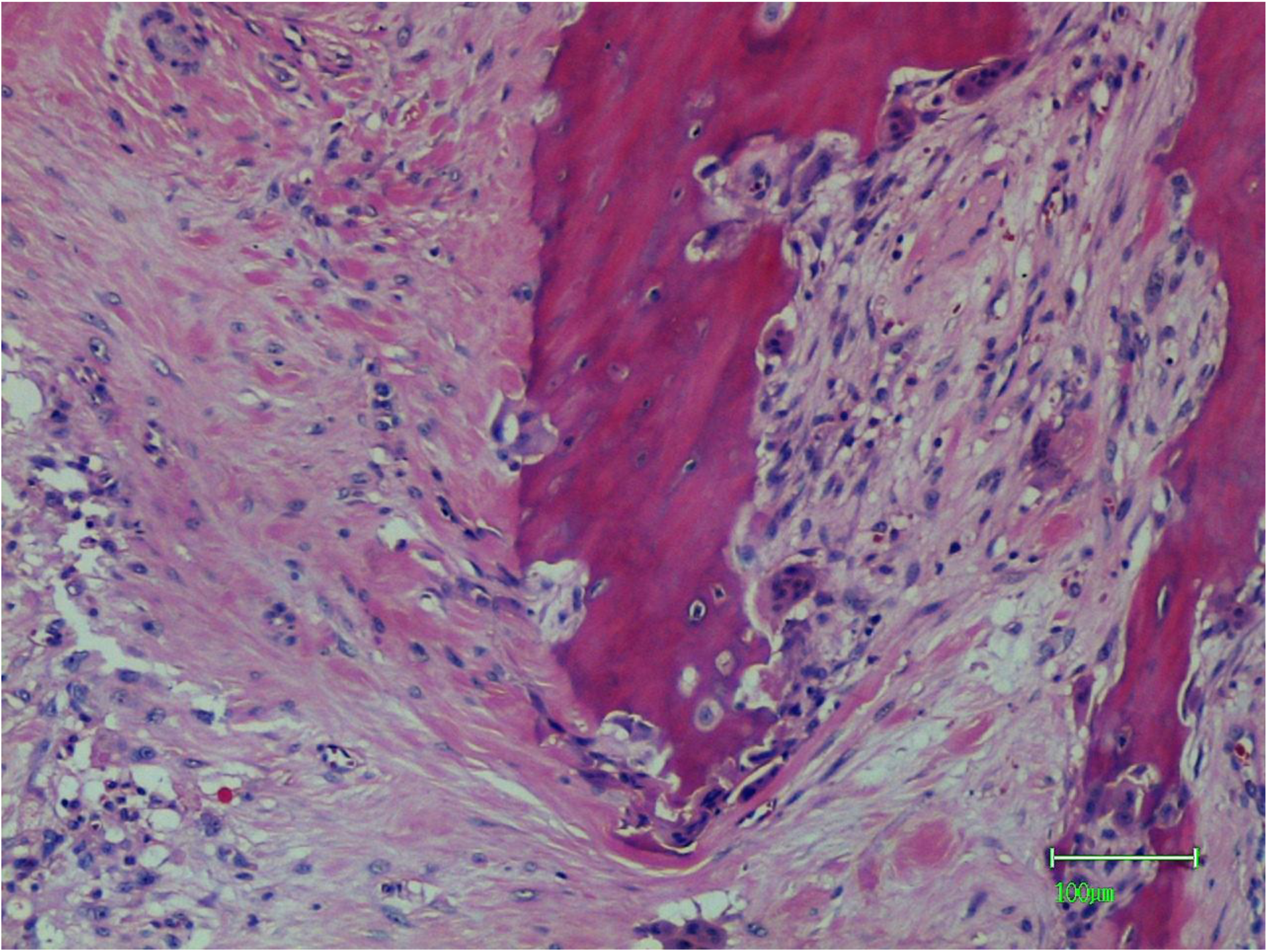
Resorption of trabecular bone by osteoclasts. Tumour is present in the bottom left corner and is separated from the bone structures by fibrous connective tissue (HE).

**Figure 9.**
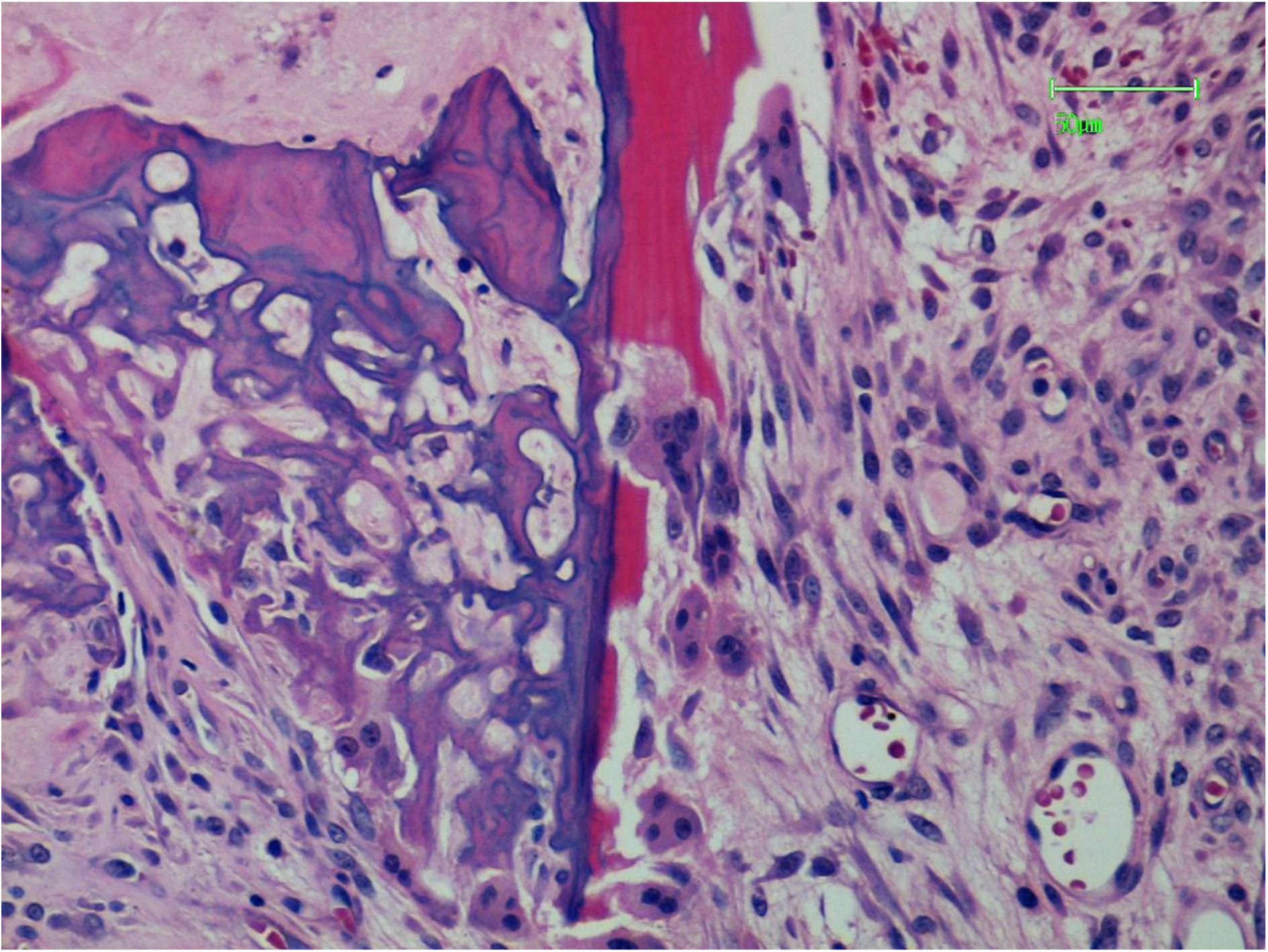
To the right there is original eosinophilic (red) bone matrix undergoing resorption by osteoclasts which are backed by a vascular mesenchymal tissue. To the left there is adherent basophilic (blue) calcified sarcoma matrix within which are several active osteoclasts (HE).

**Figure 10.**
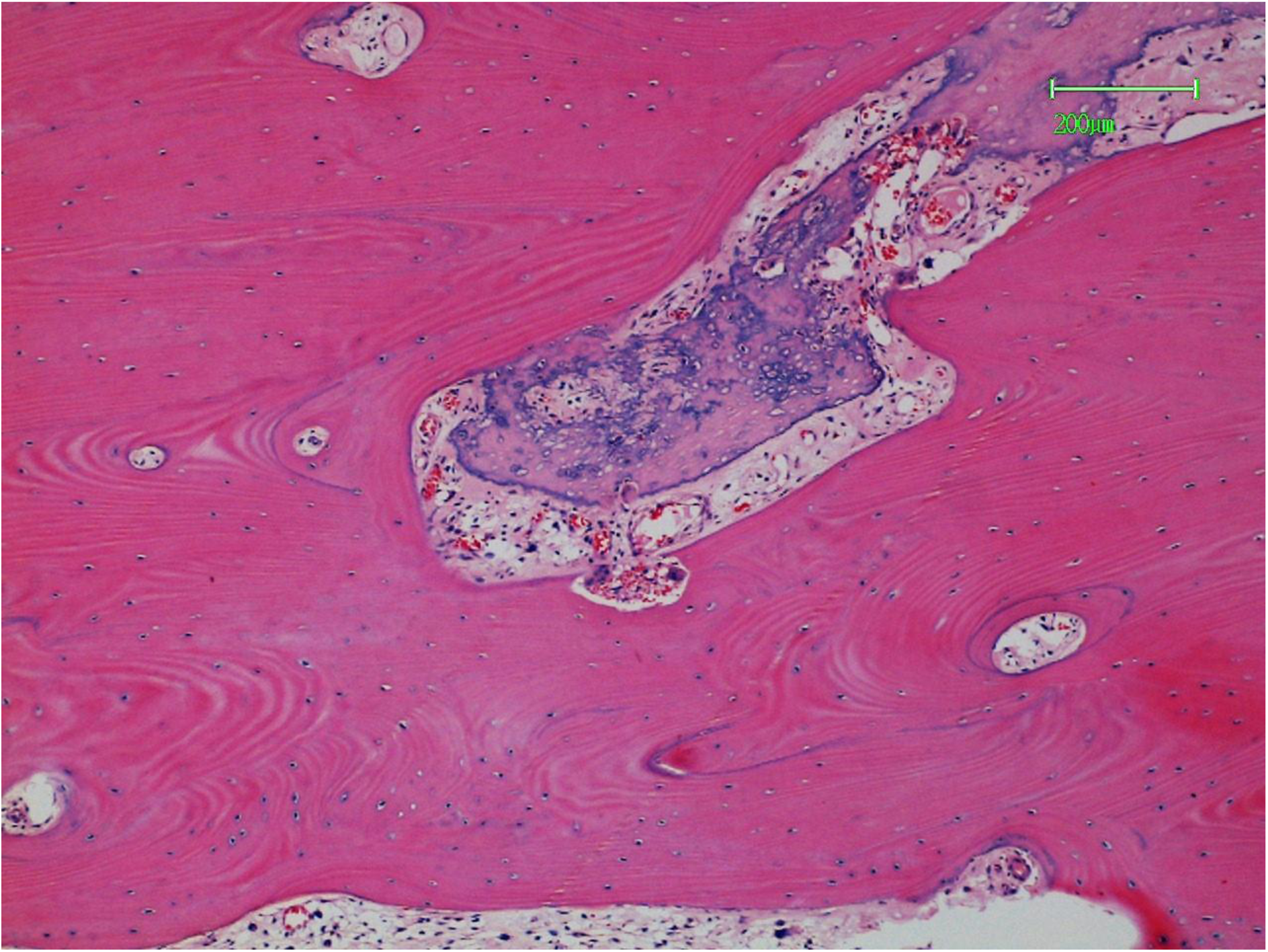
Sarcoma (basophilic matrix) has permeated a vascularized channel in cortical bone (HE).

**Figure 11.**
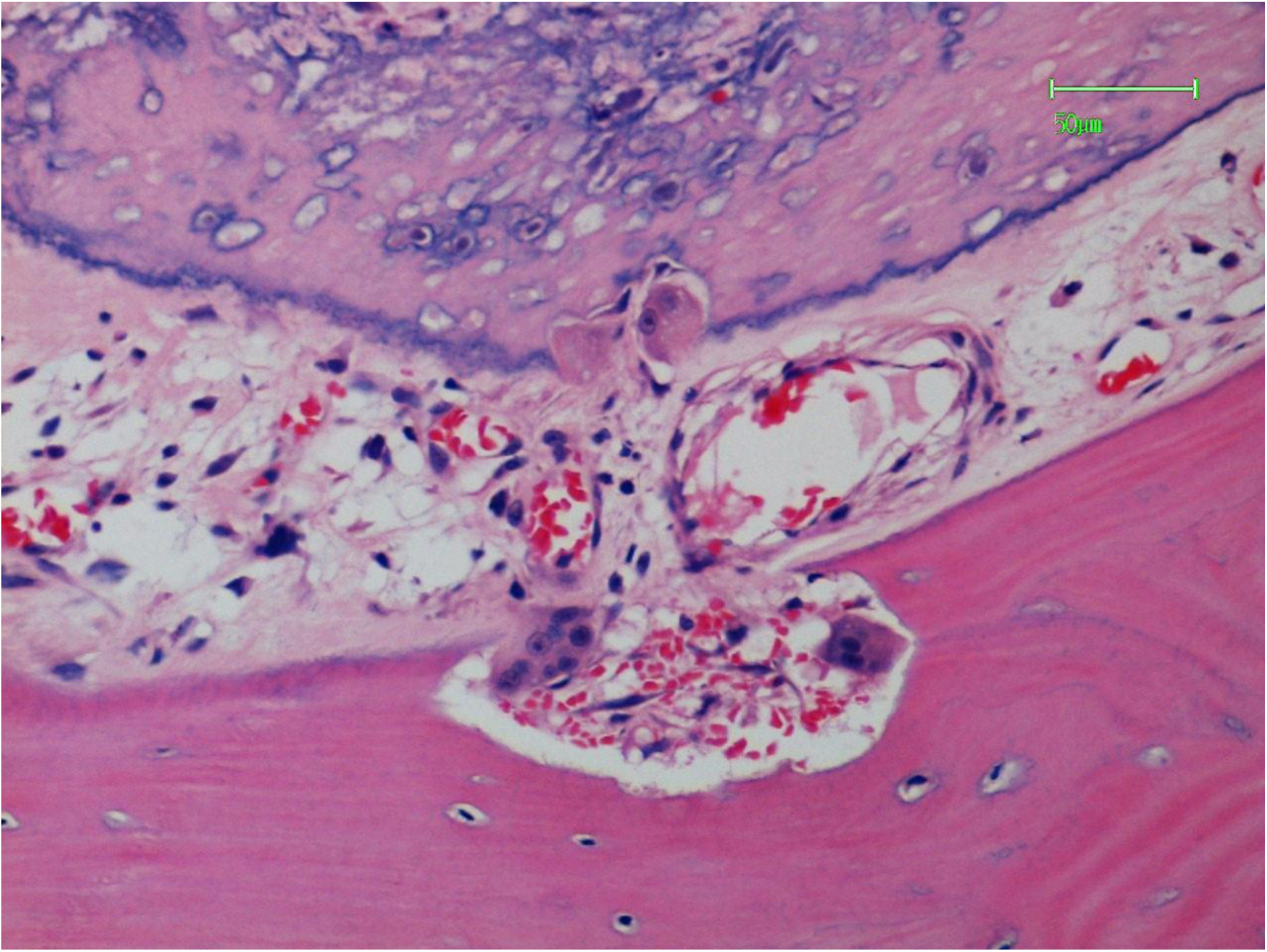
Higher power view of sample in Figure 10. Towards the six o’clock position osteoclasts and companion blood vessels are cutting into cortical bone. Towards the 12 o’clock position there is an osteoclast resorbing tumour bone (HE).

**Figure 12.**
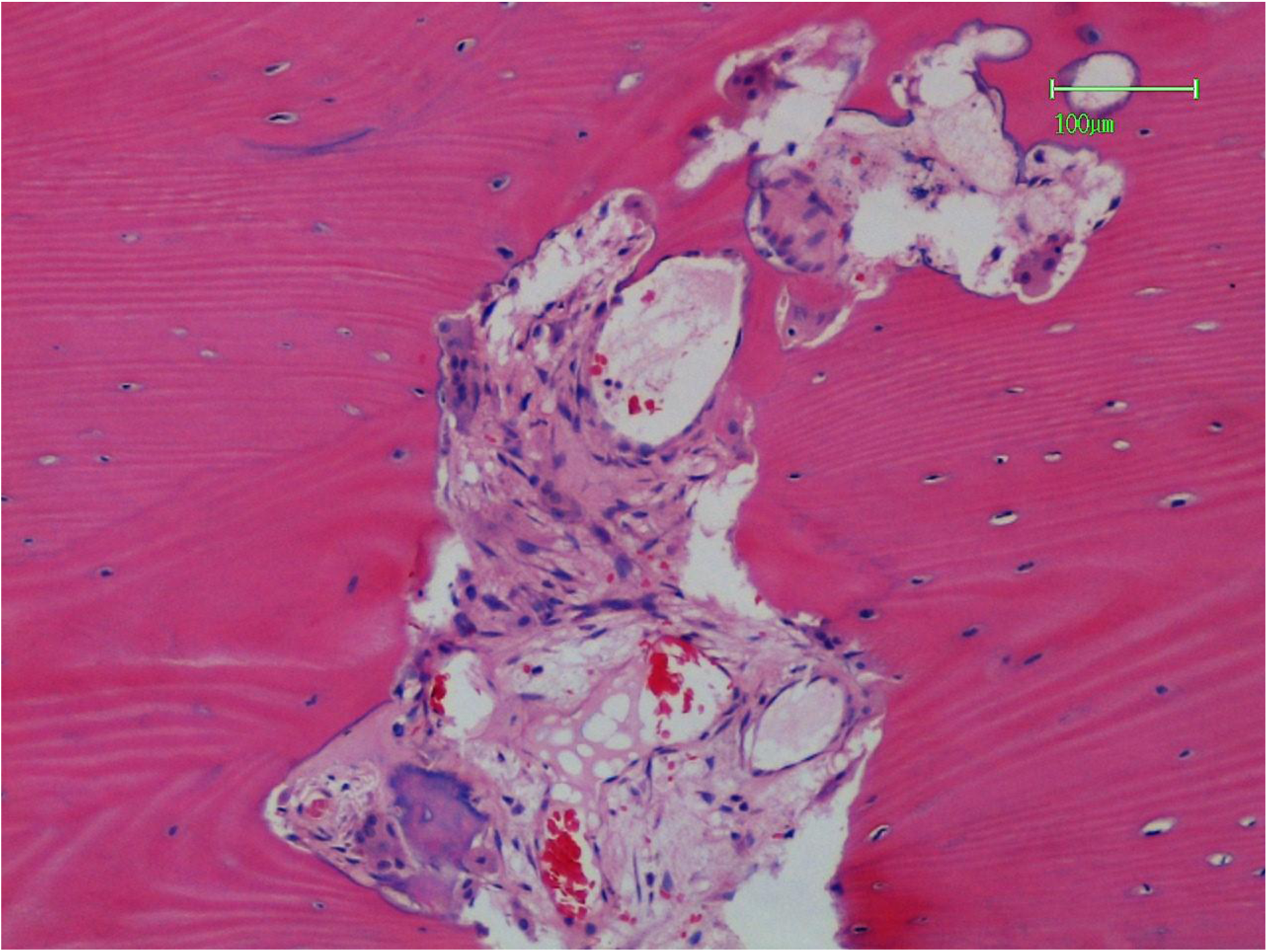
Osteoclasts cutting a vascularized channel in cortical bone creating a complex two-dimensional pattern. Some tumour bone is present in the 7 o’clock position with an attendant osteoclast (HE).

**Figure 13.**
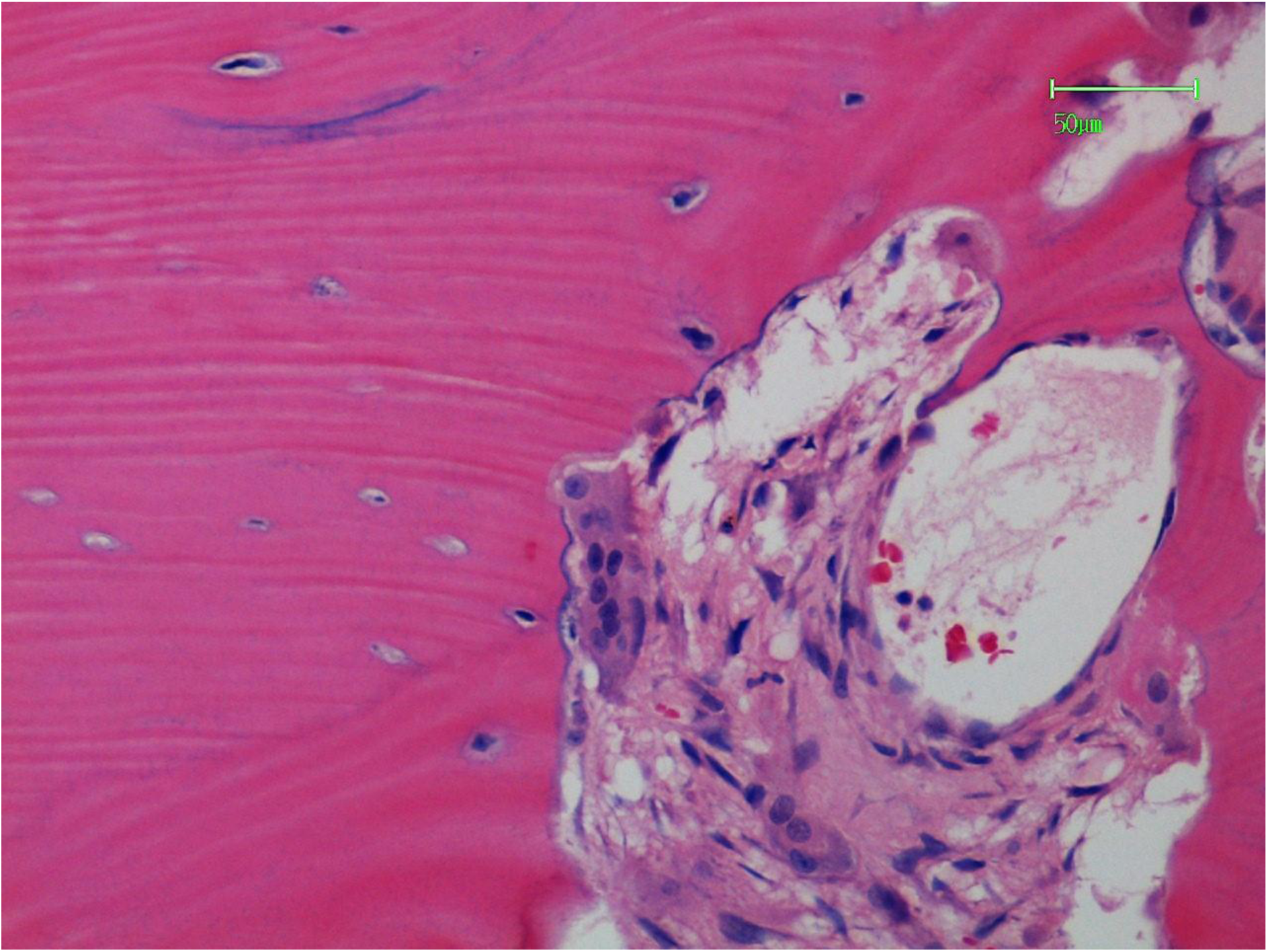
A higher power view of the field shown in Figure 12. The osteoclast is cutting at right angles to the lamellae of the cortical bone (HE).

**Figure 14.**
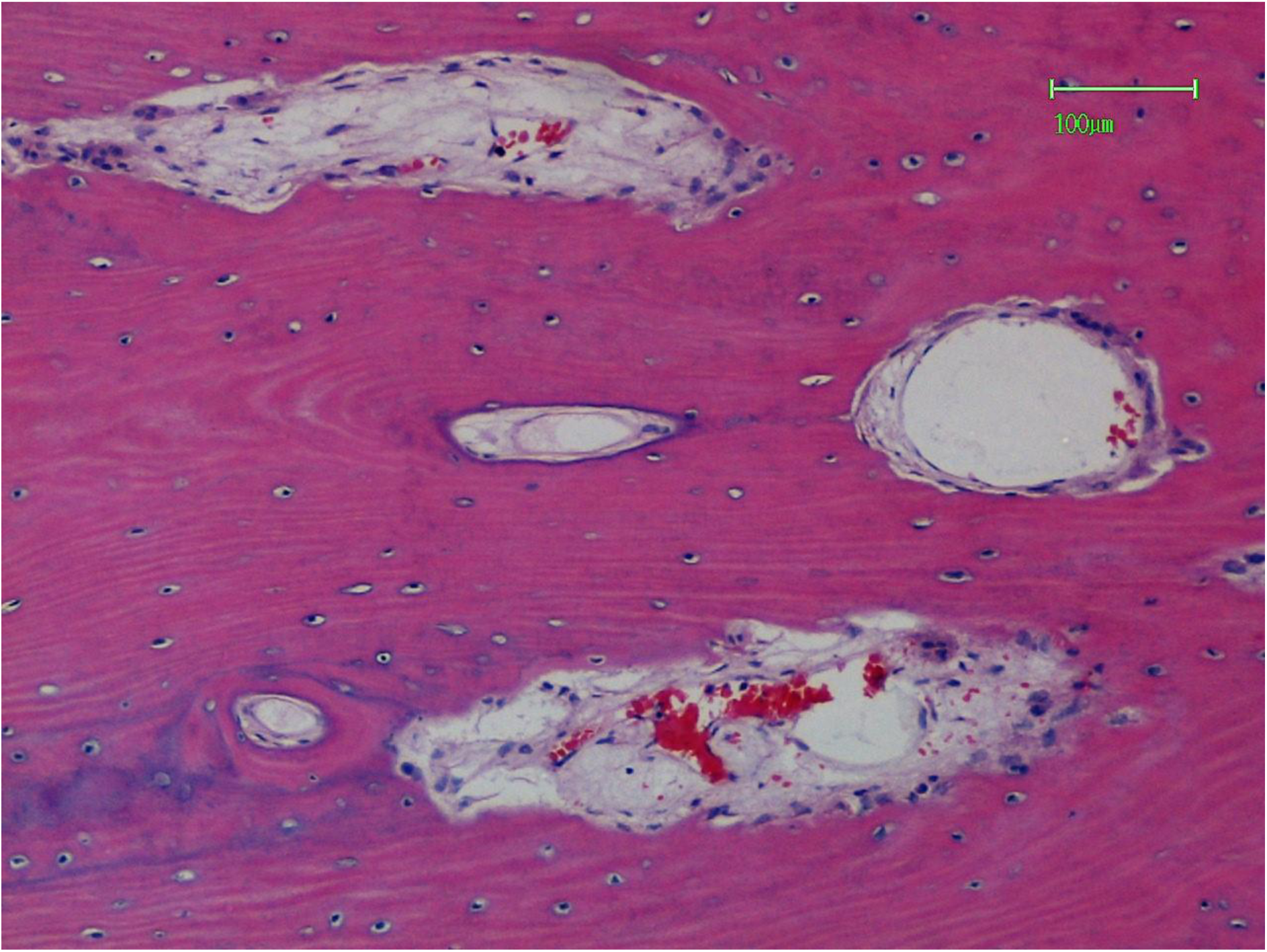
Three osteoclast cutting channels are present together with two Haversian canals in cortical osteonal bone. The lower (and to the left) Haversian canal is closely related to the lowest cutting channel (HE).

**Figure 15.**
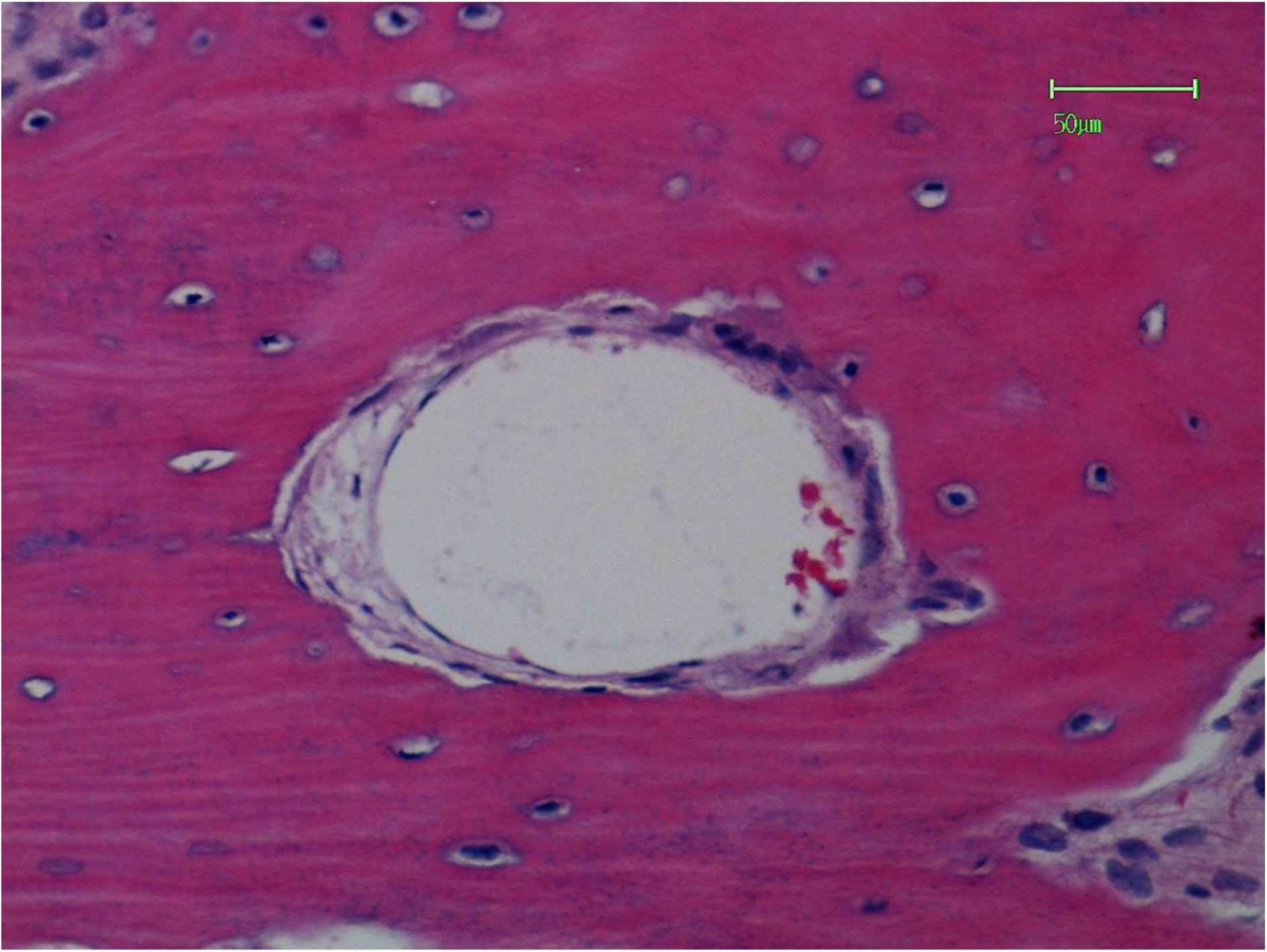
A cutting channel from Figure 14 showing osteoclasts and a markedly dilated capillary (HE).

**Figure 16.**
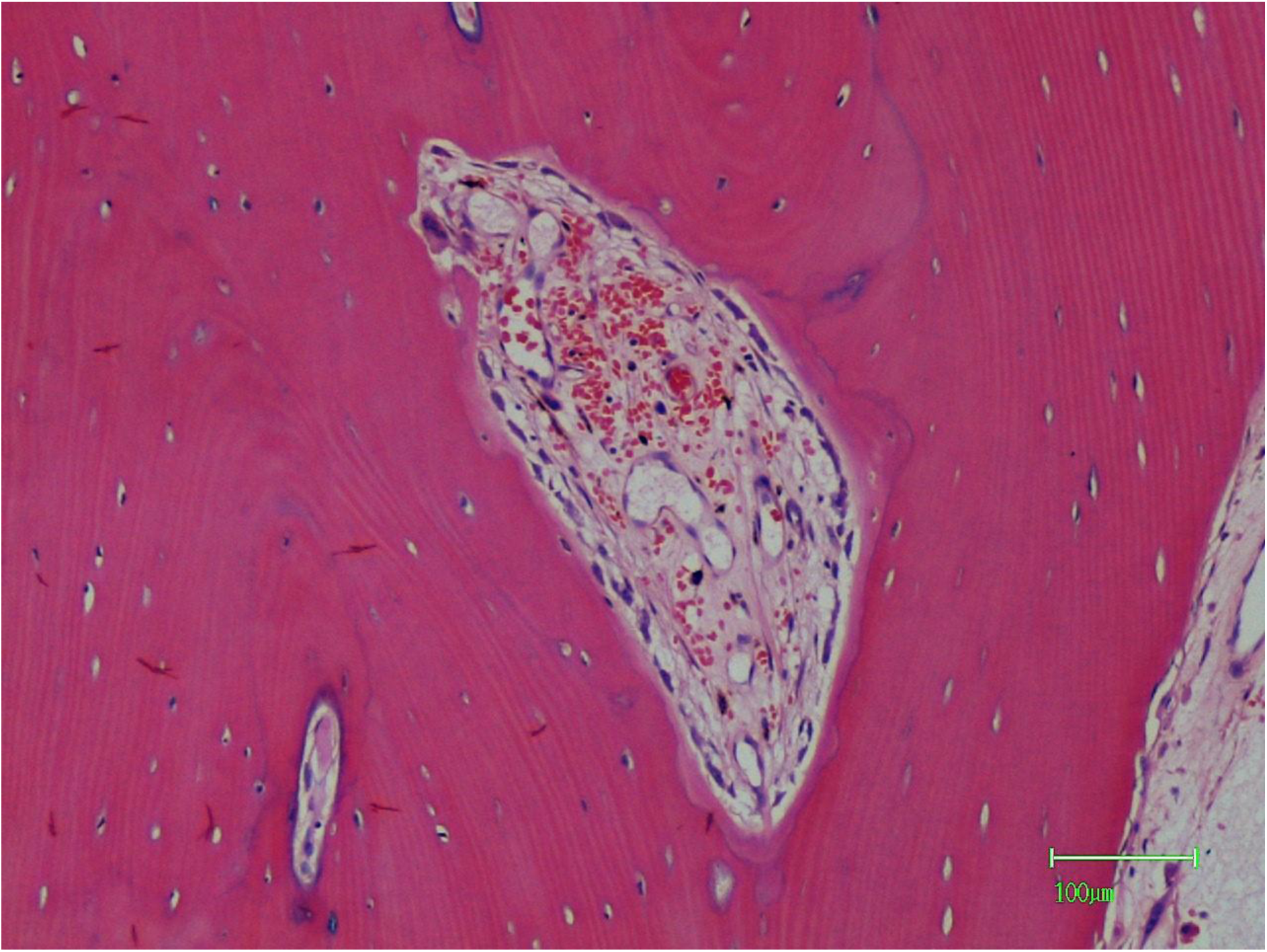
A vascularized cutting channel with a single osteoclast at the 11 o’clock position. The wall of the channel is lined by osteoblasts which have formed a thin layer of new bone. Tumour is not present in this channel. A Haversian canal is at the lower left (HE).

**Figure 17.**
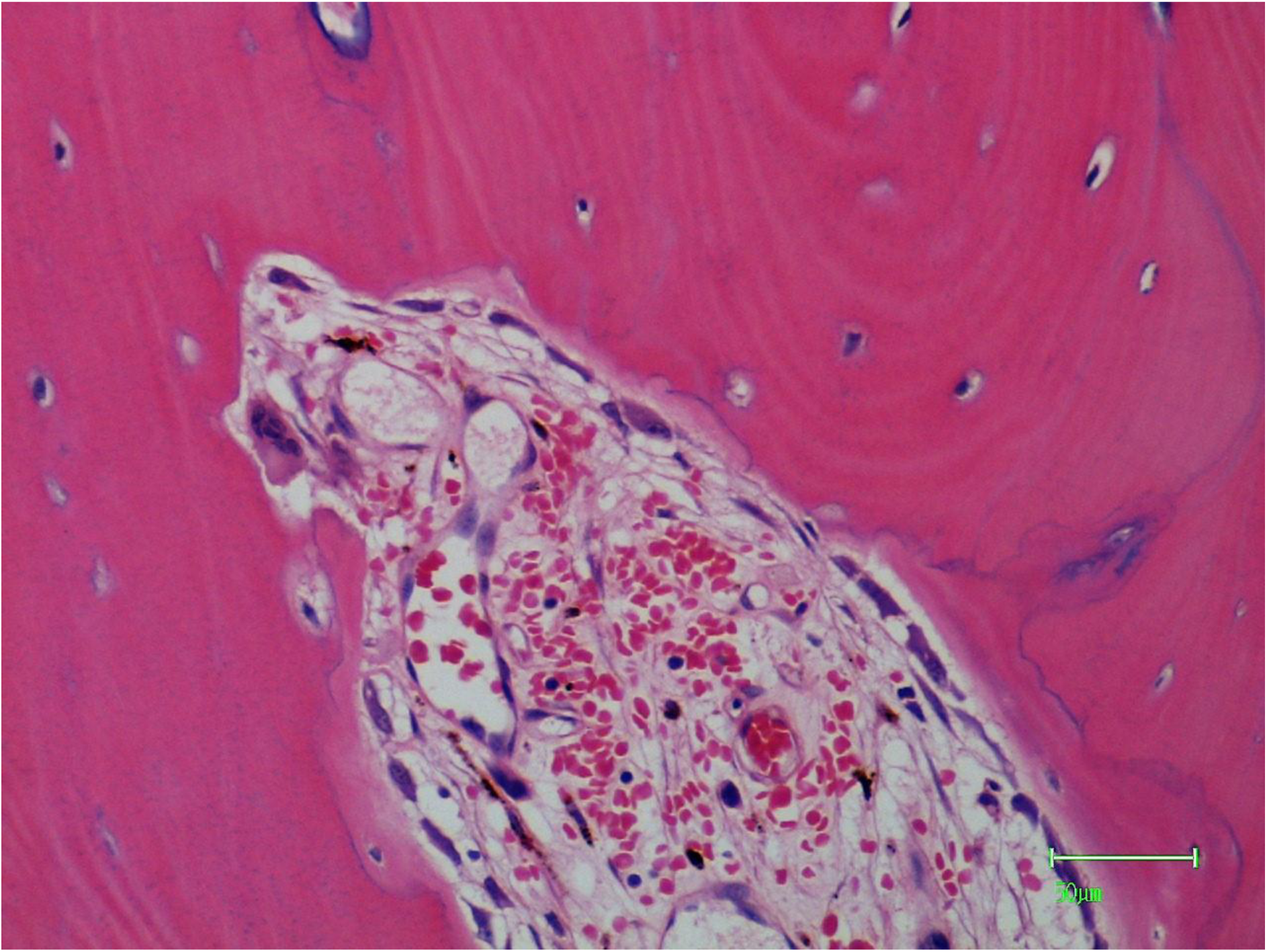
A higher power view of part of Figure 16. The osteoclast is closely related to a capillary. Mast cells are also present (HE).

**Figure 18.**
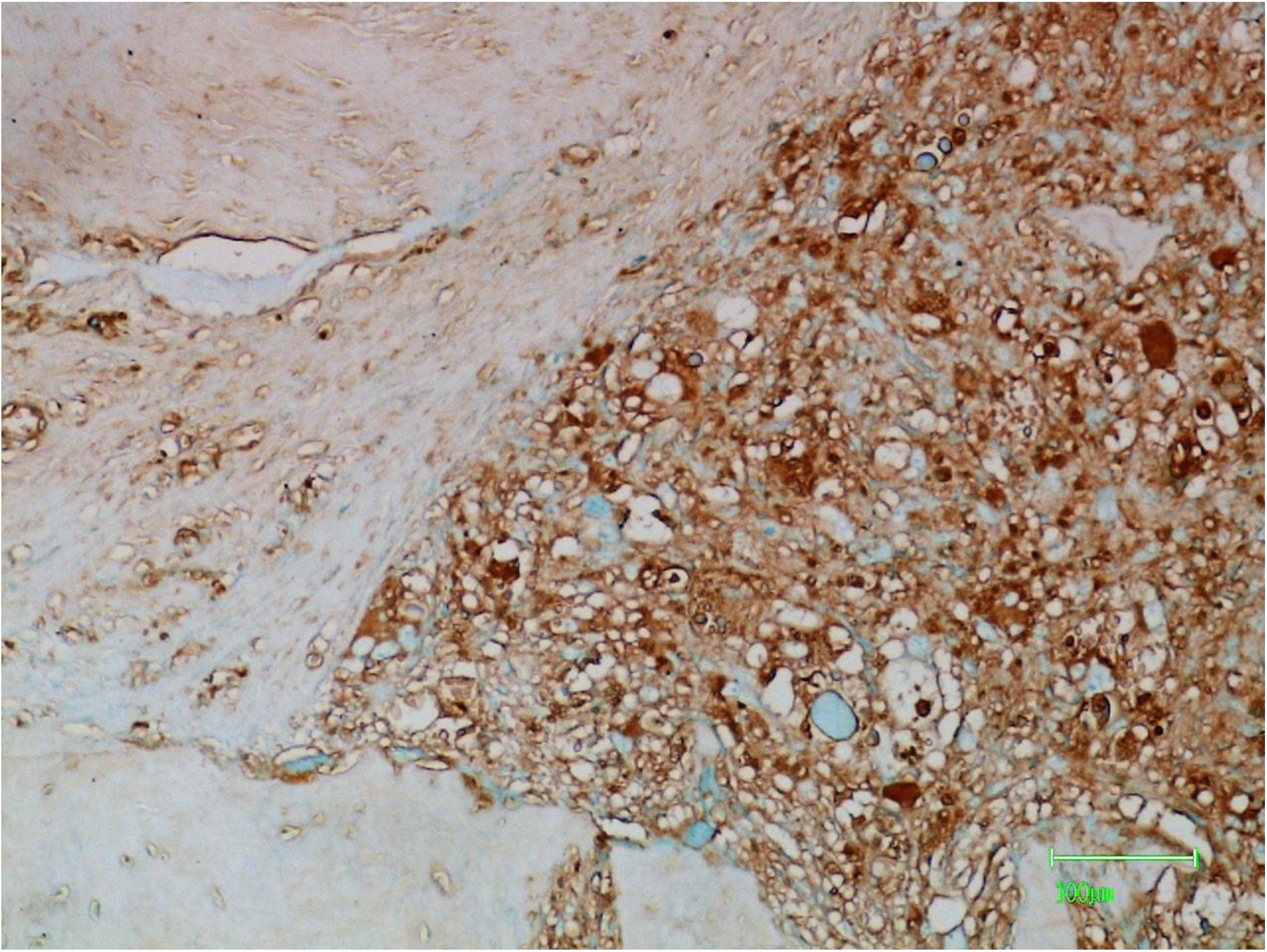
Osteogenic sarcoma showing cytoplasmic and membrane positive reactions with l-PHA (brown staining).

**Figure 19.**
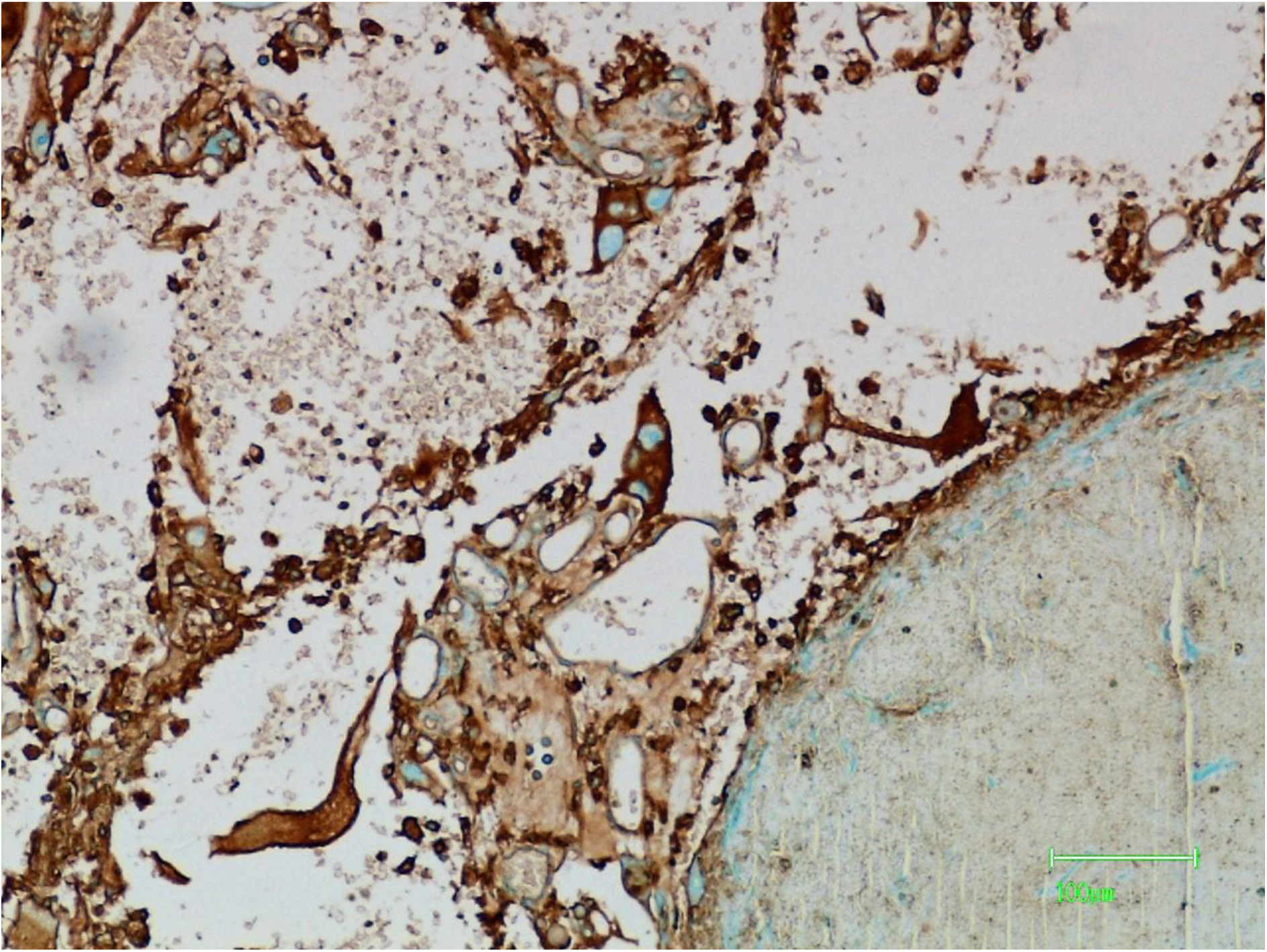
Pleomorphic tumour cells showing positive reactions with l-PHA.

**Figure 20.**
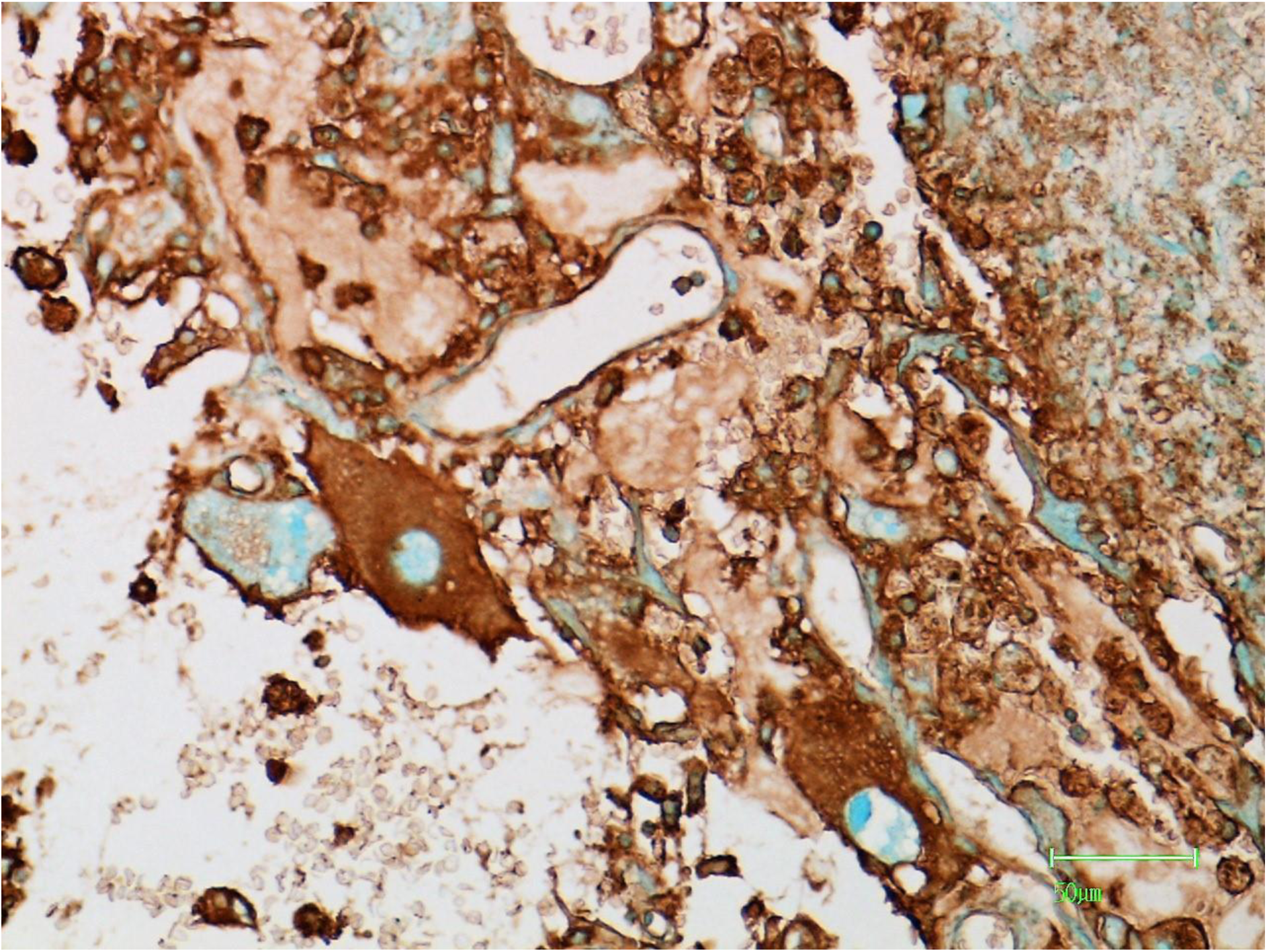
Positive staining with l-PHA in pleomorphic sarcoma cells and flattened capillary endothelial cells.

**Figure 21.**
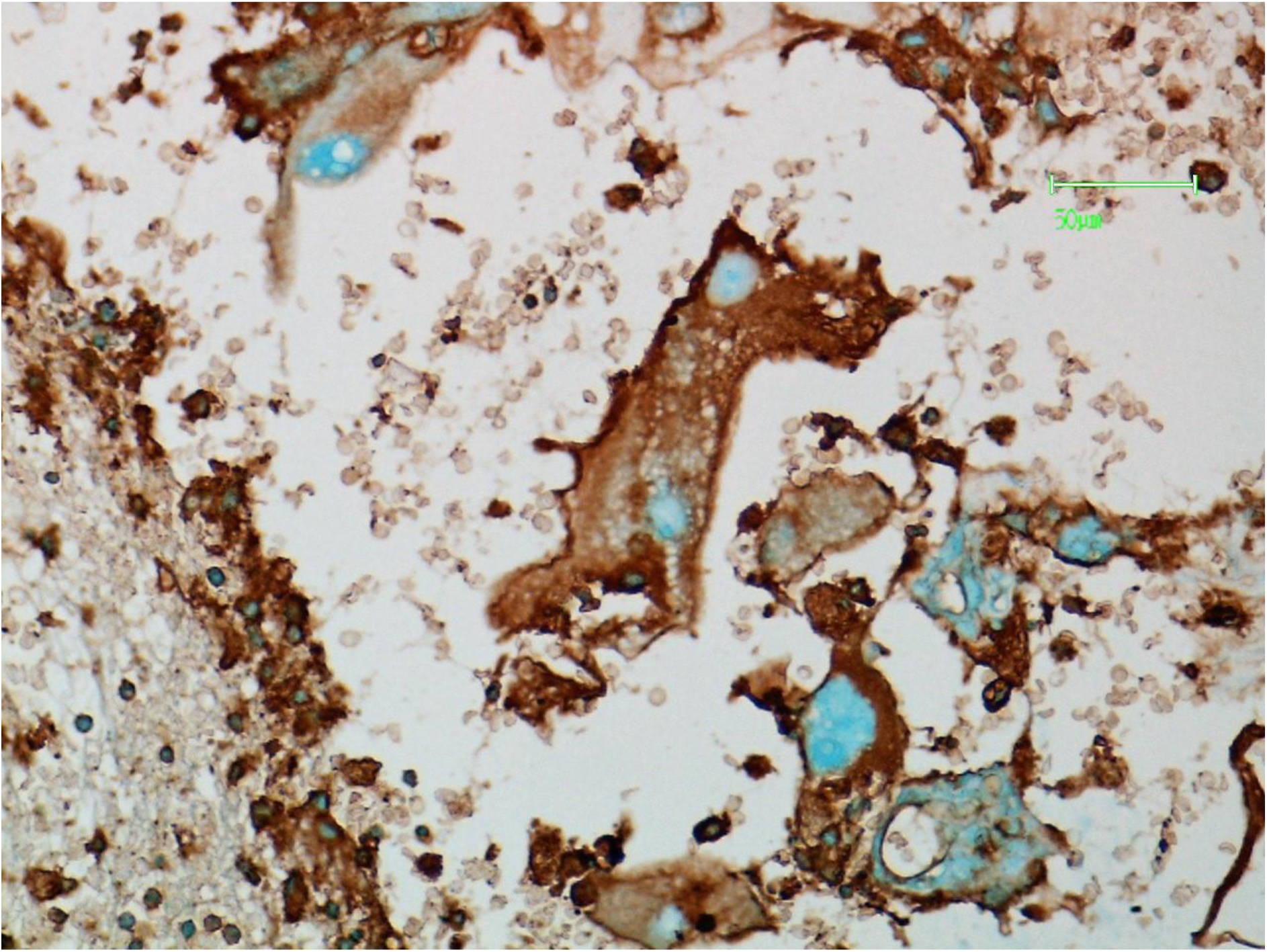
Sarcoma cells showing positive staining reactions with l-PHA. Intranuclear reactions are negative.

**Figure 22.**
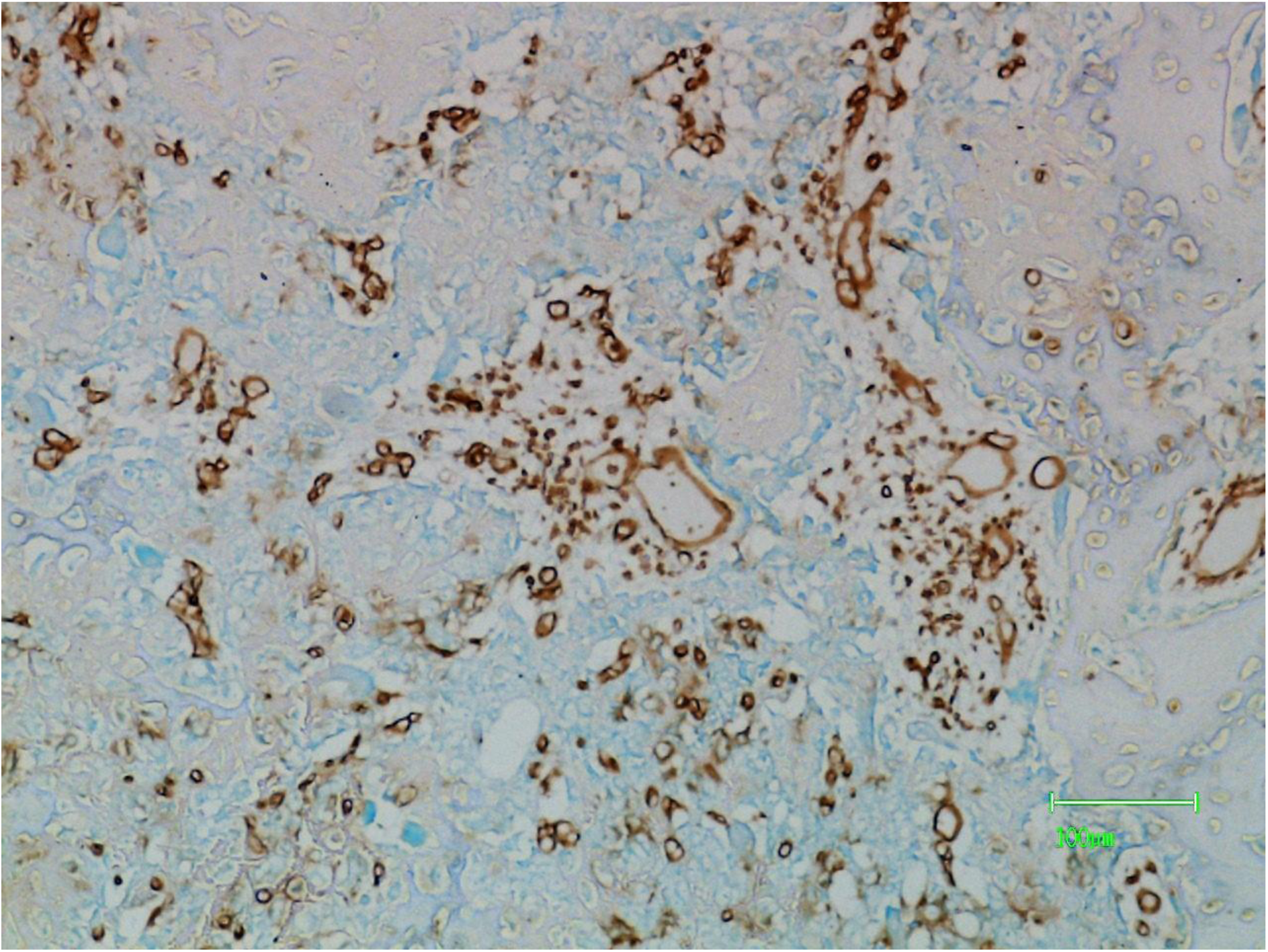
Intratumoural vascularity is demonstrated by positive reactions with PTL-II. There is variation in vessel size and shape.

**Figure 23.**
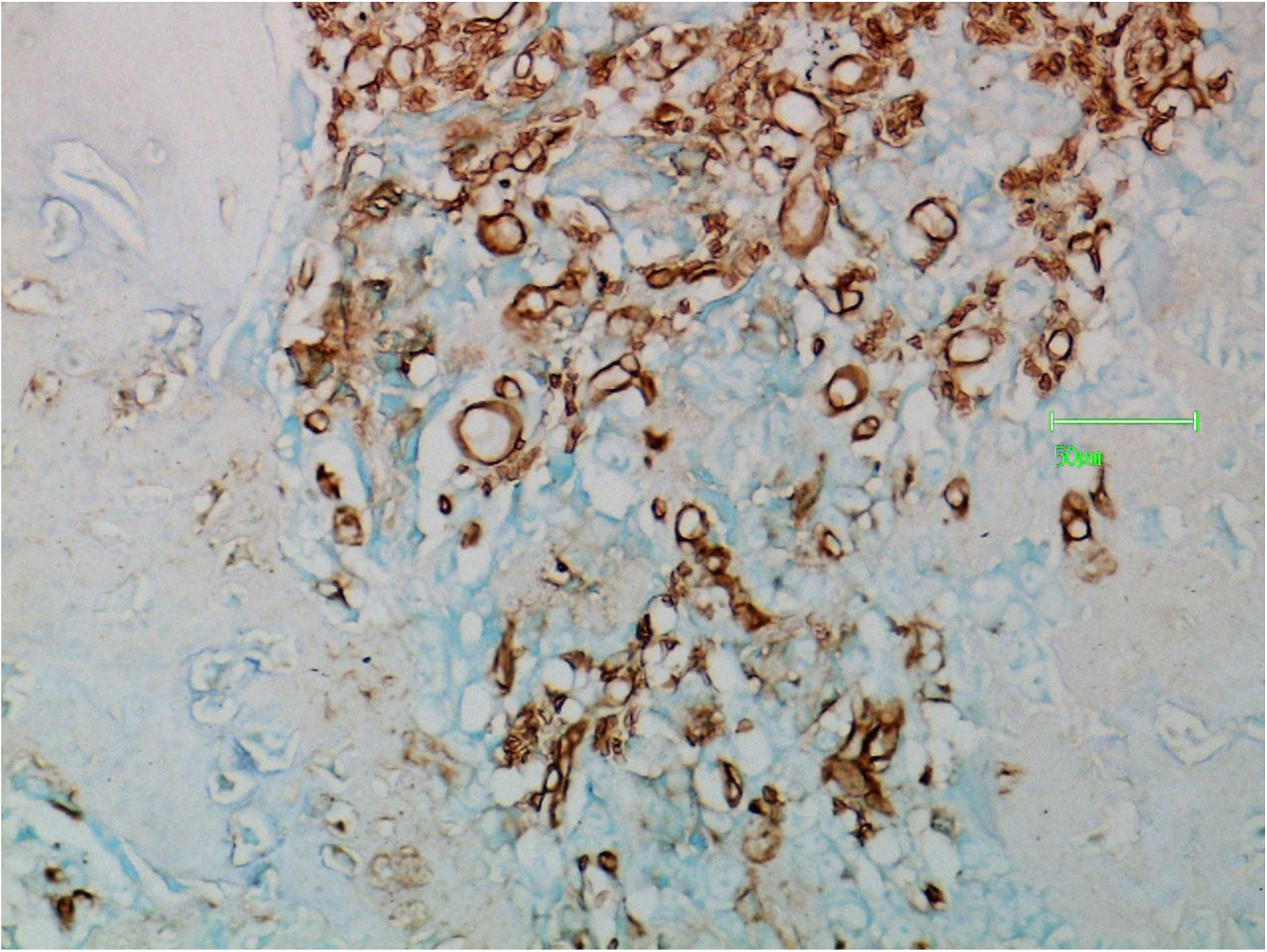
A higher power view of part of figure 22 again stained with PTL-II and again showing variation in vessel size and shape.

**Figure 24.**
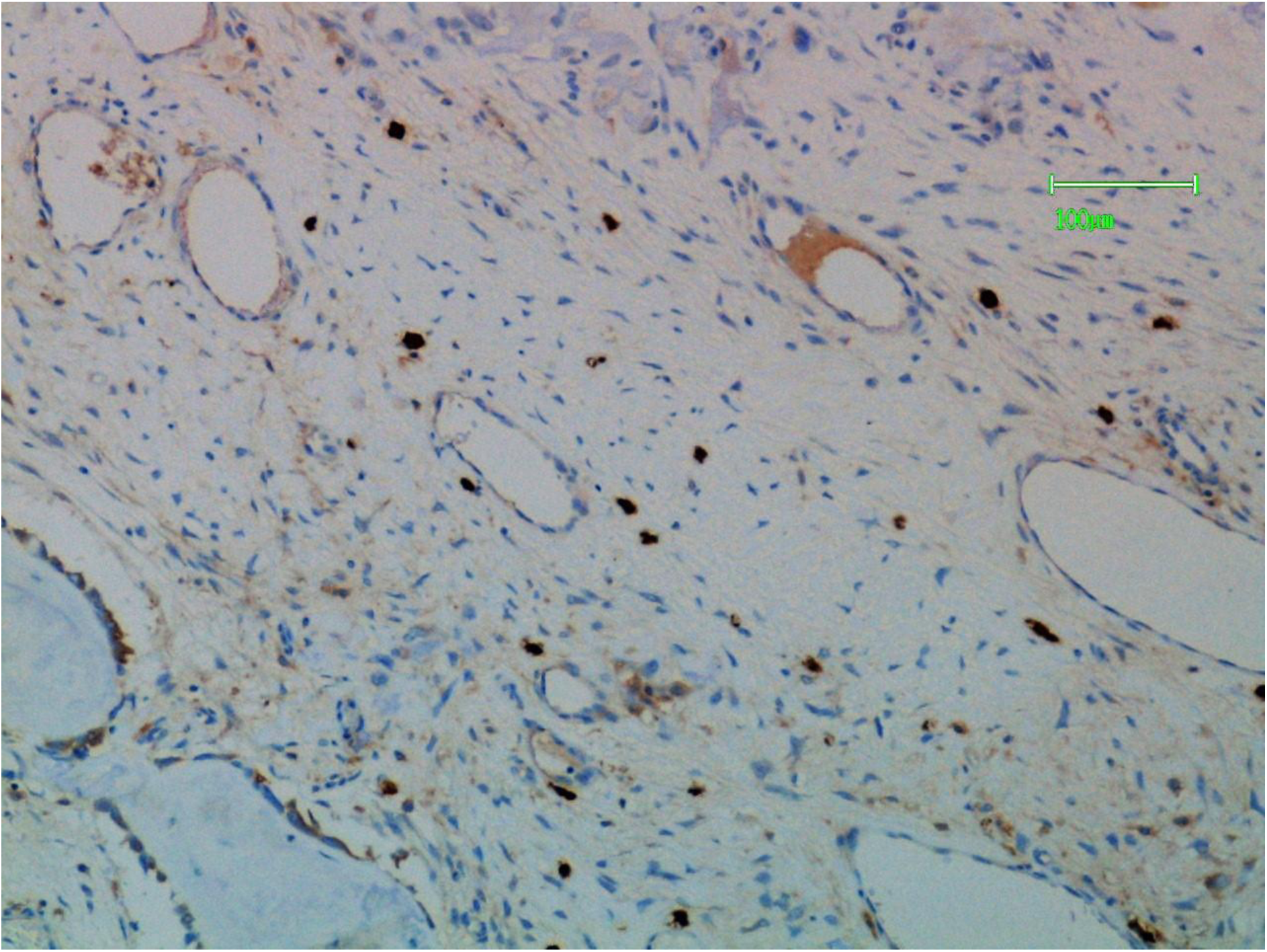
Mast cells (stained for mast cell tryptase) in connective tissue adjacent to sarcoma (top right).

**Figure 25.**
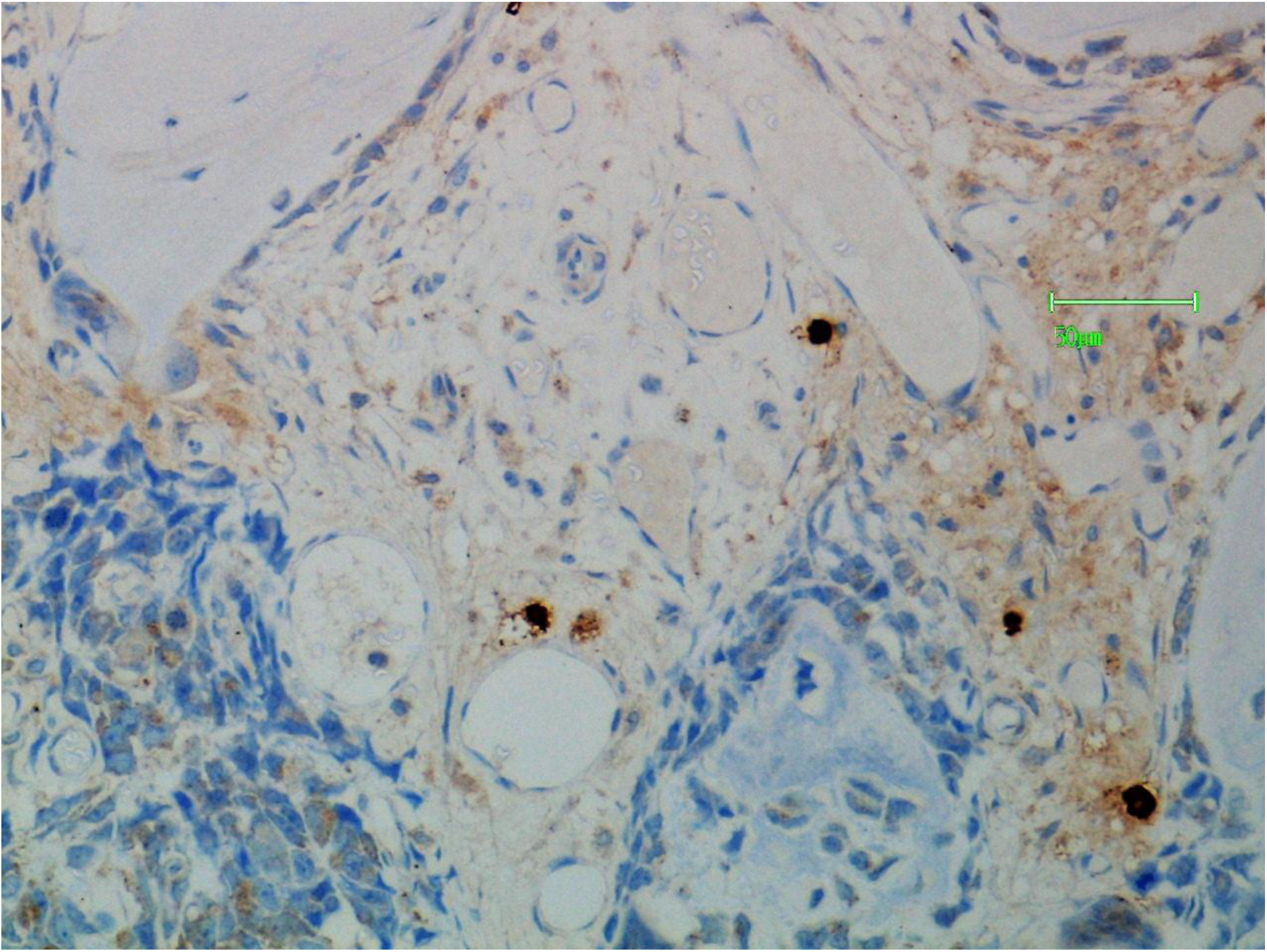
Mast cells (tryptase positive) showing degranulation in relation to sarcoma (bottom of field of view).

**Figure 26.**
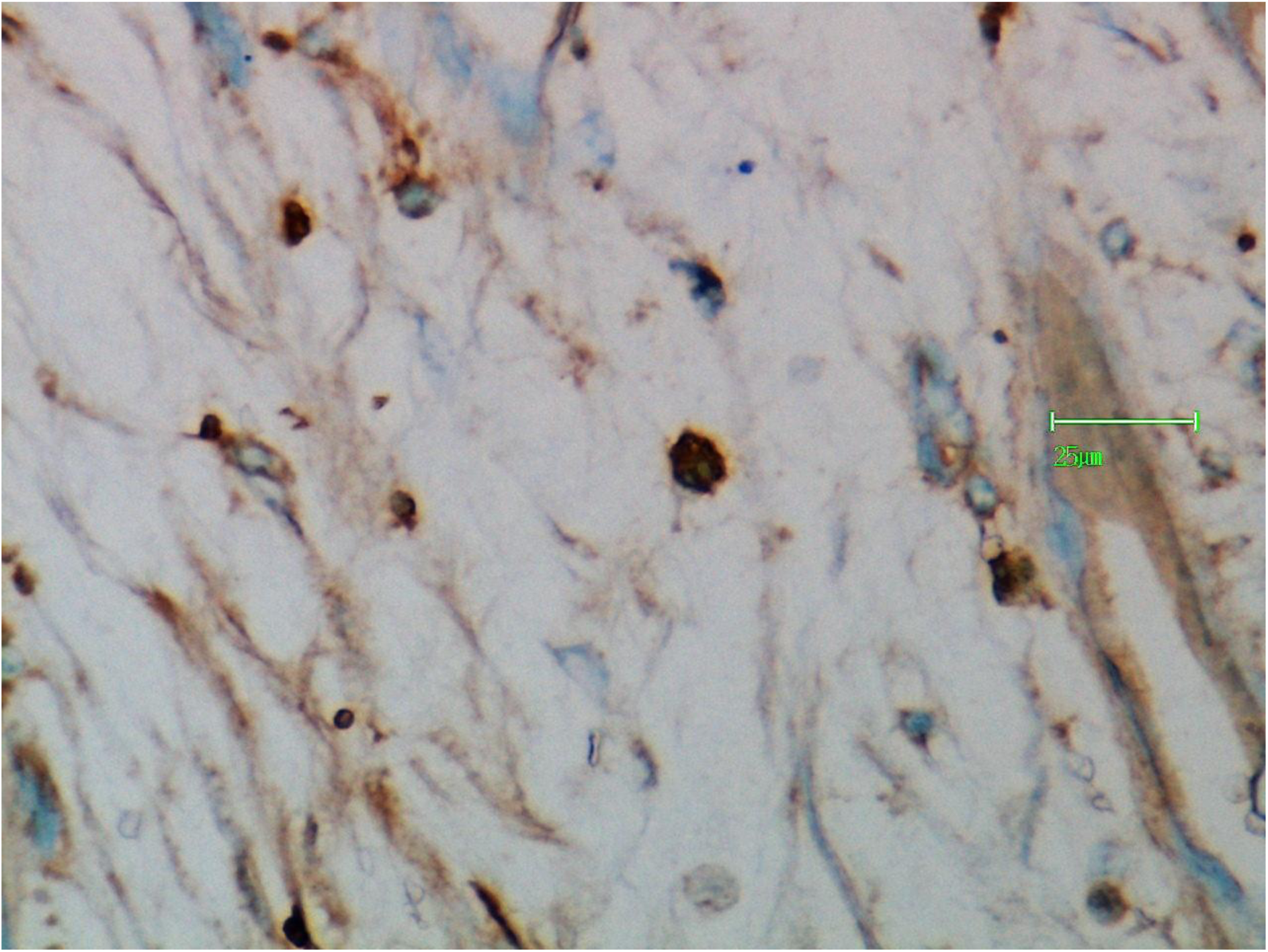
Mast cells showing a positive cytoplasmic reaction with l-PHA.

**Figure 27.**
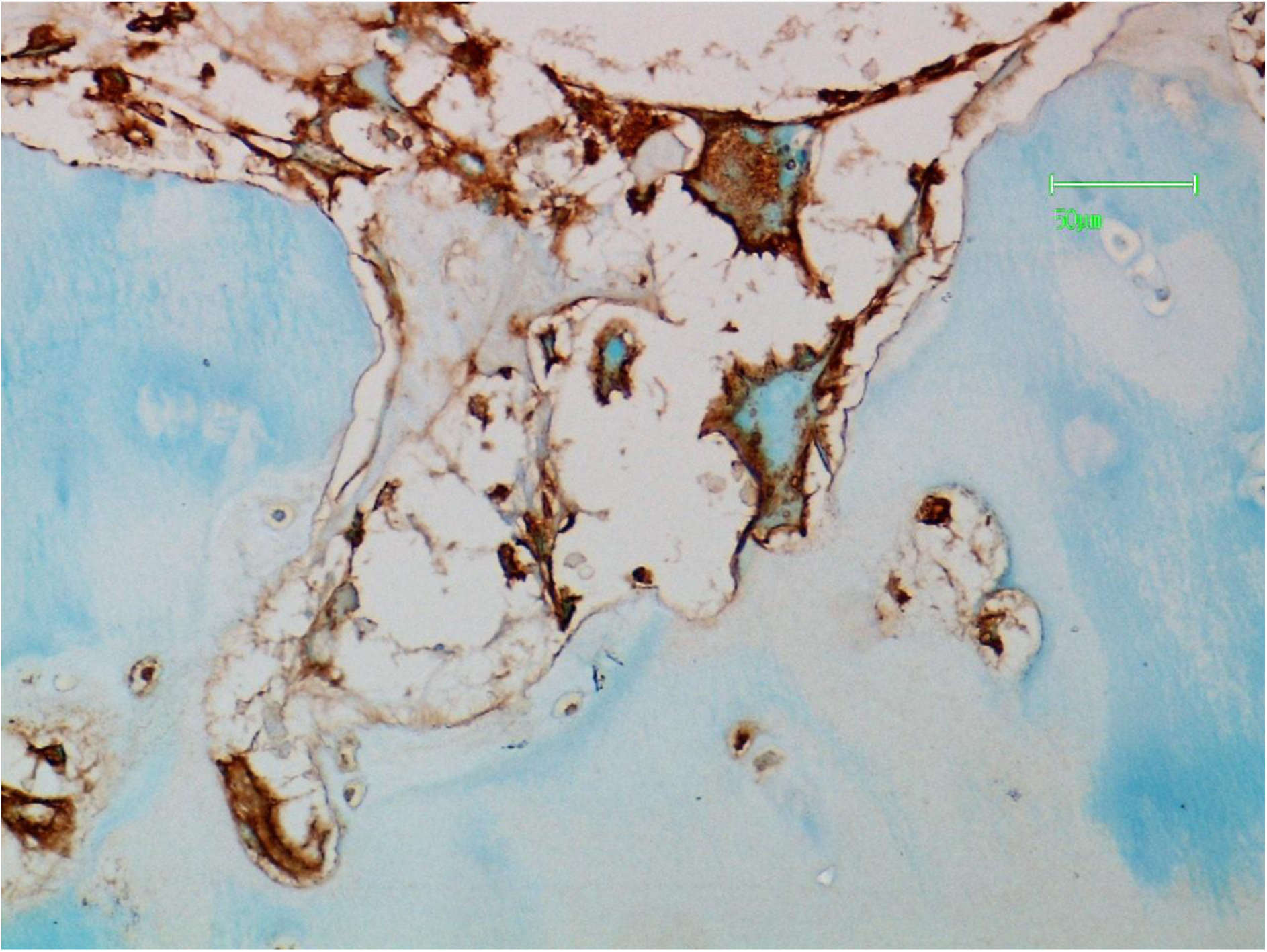
Tumour cells (large and pleomorphic) showing positive reaction with l-PHA. An osteoclast at seven o’clock also shows positive reaction both cytoplasmic and membranous in location but accentuated in the latter.

**Figure 28.**
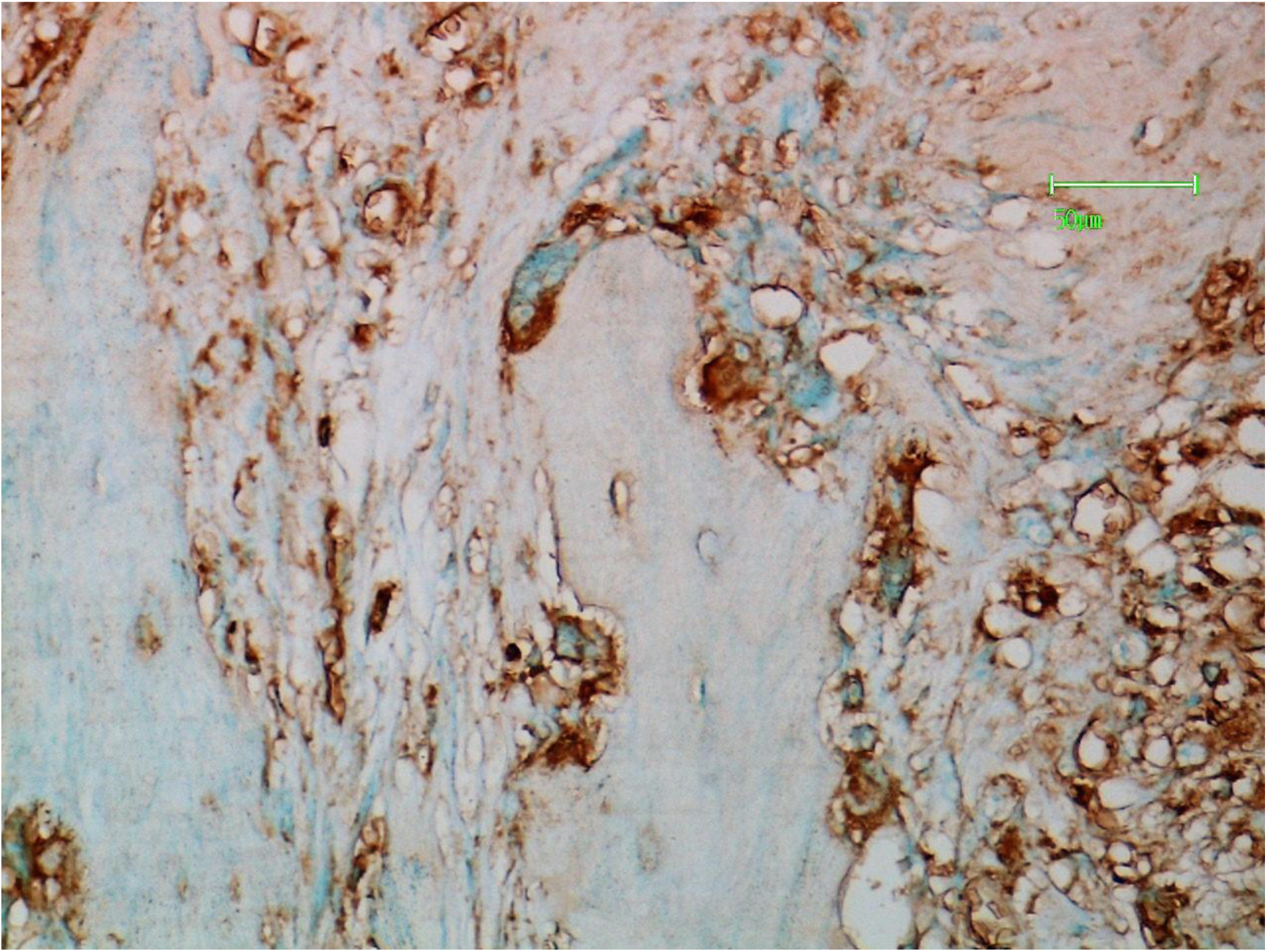
Osteoclasts and capillary endothelial cells are positive with l-PHA. The reaction is most prominent on the resorptive surface.

**Figure 29.**
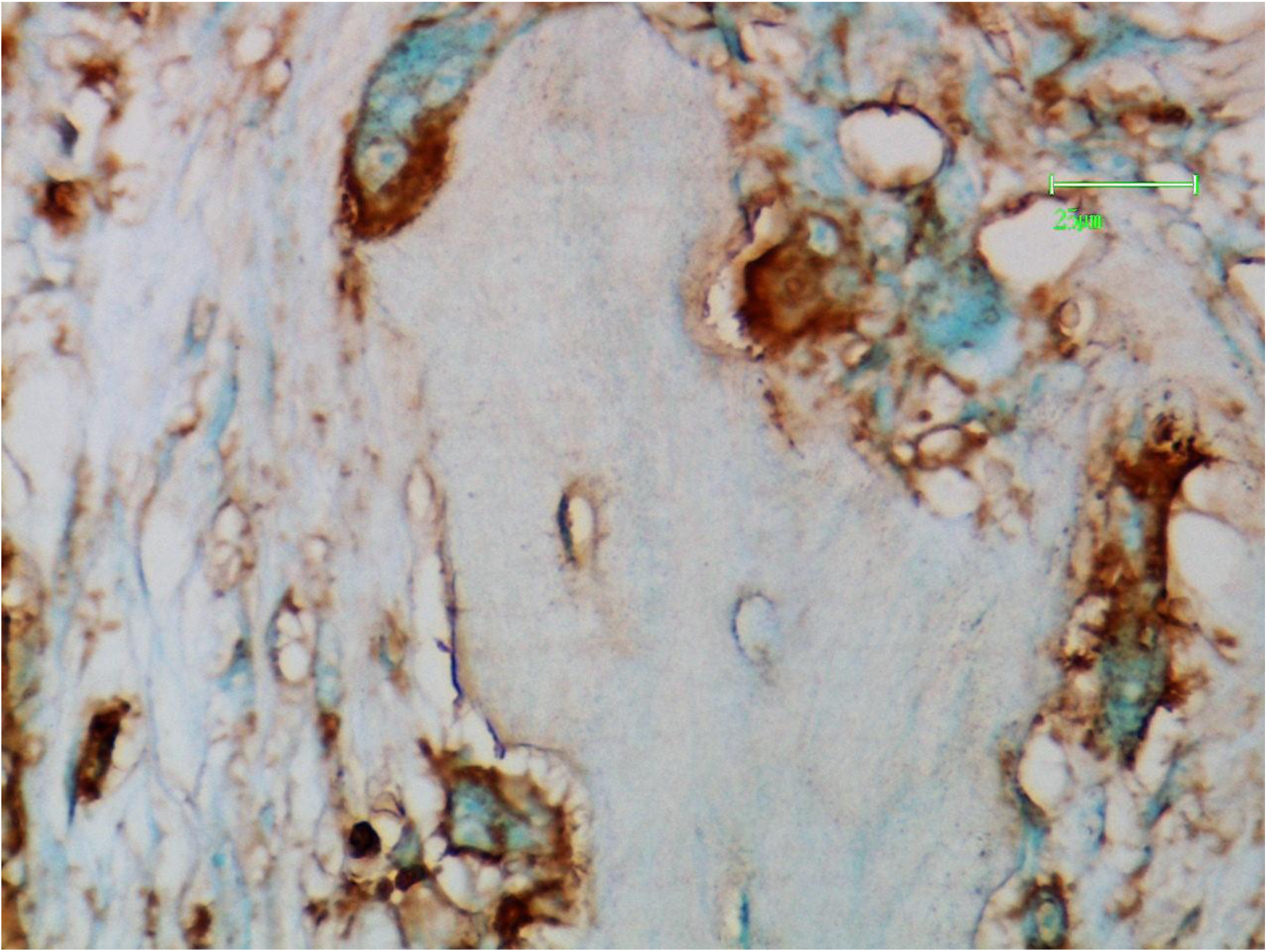
Higher power view emphasising the resorptive zone of positivity of osteoclasts with l-PHA.

**Figure 30.**
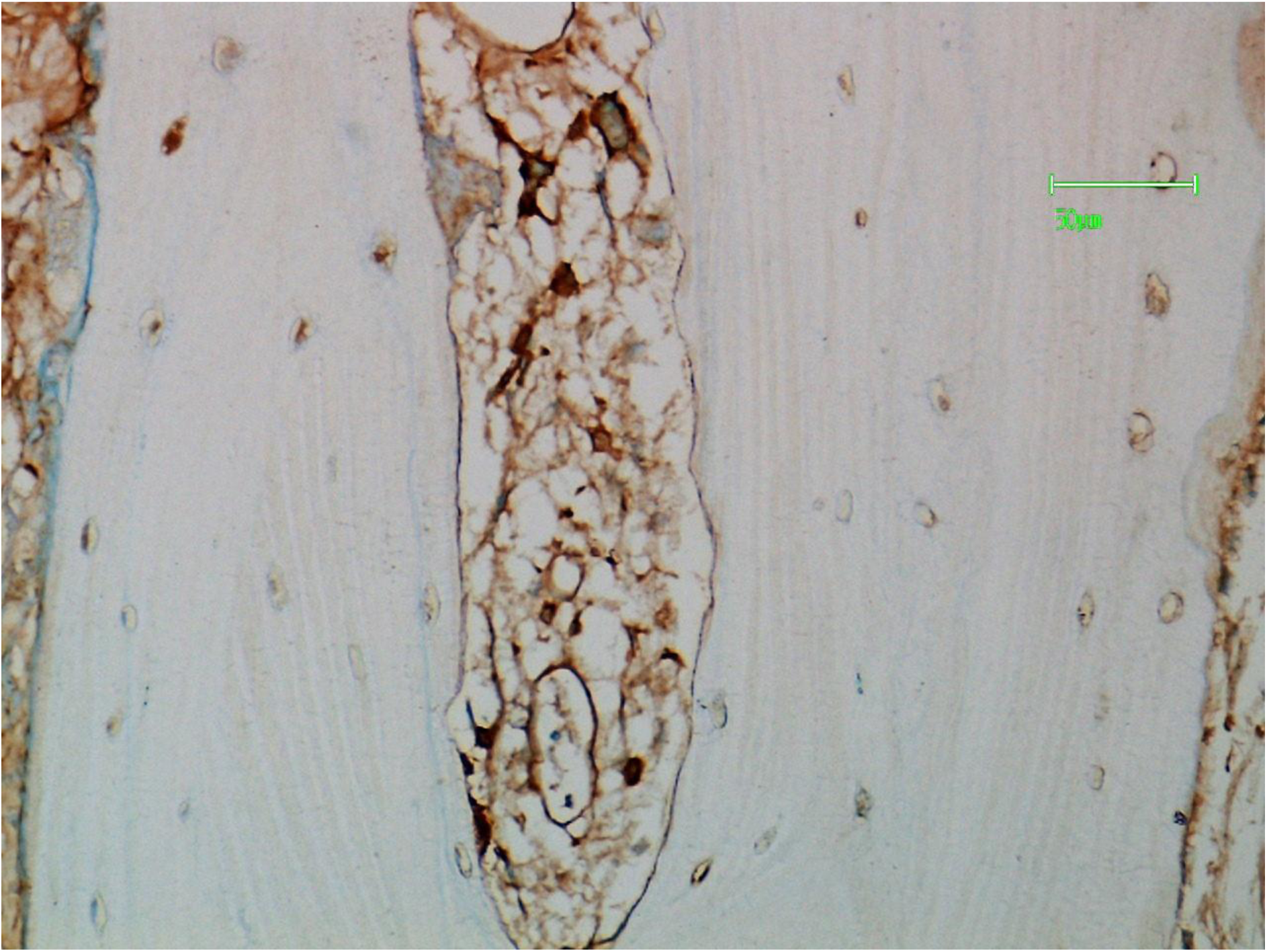
Mast cells and endothelial cells in the cutting channel in cortical bone are positive with l-PHA.

**Figure 31.**
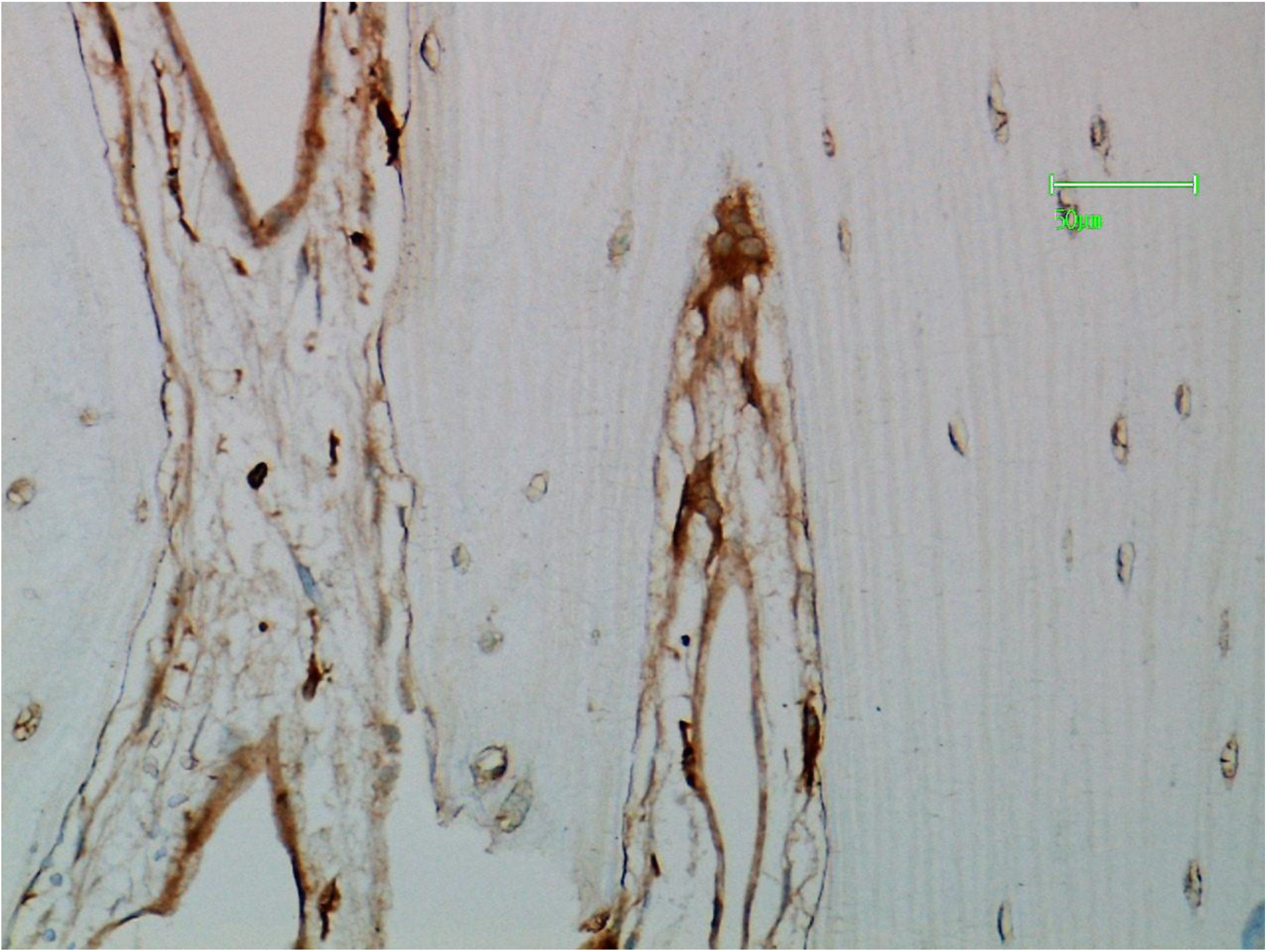
Osteoclasts in the cutting channel are positive for l-PHA.

**Figure 32.**
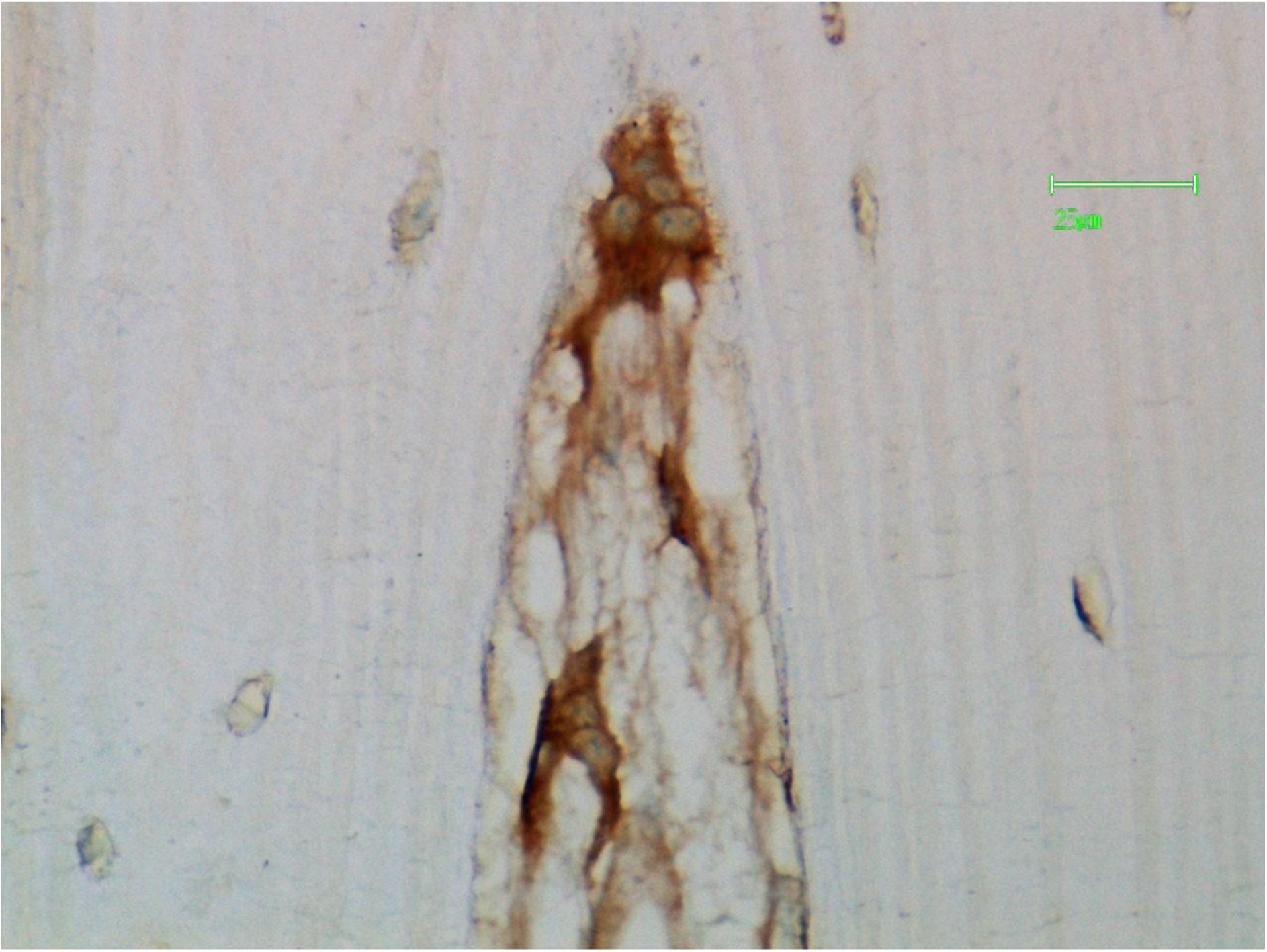
Higher power view of an osteoclast at the apex of a cutting channel confirming positivity for l-PHA.

**Figure 33.**
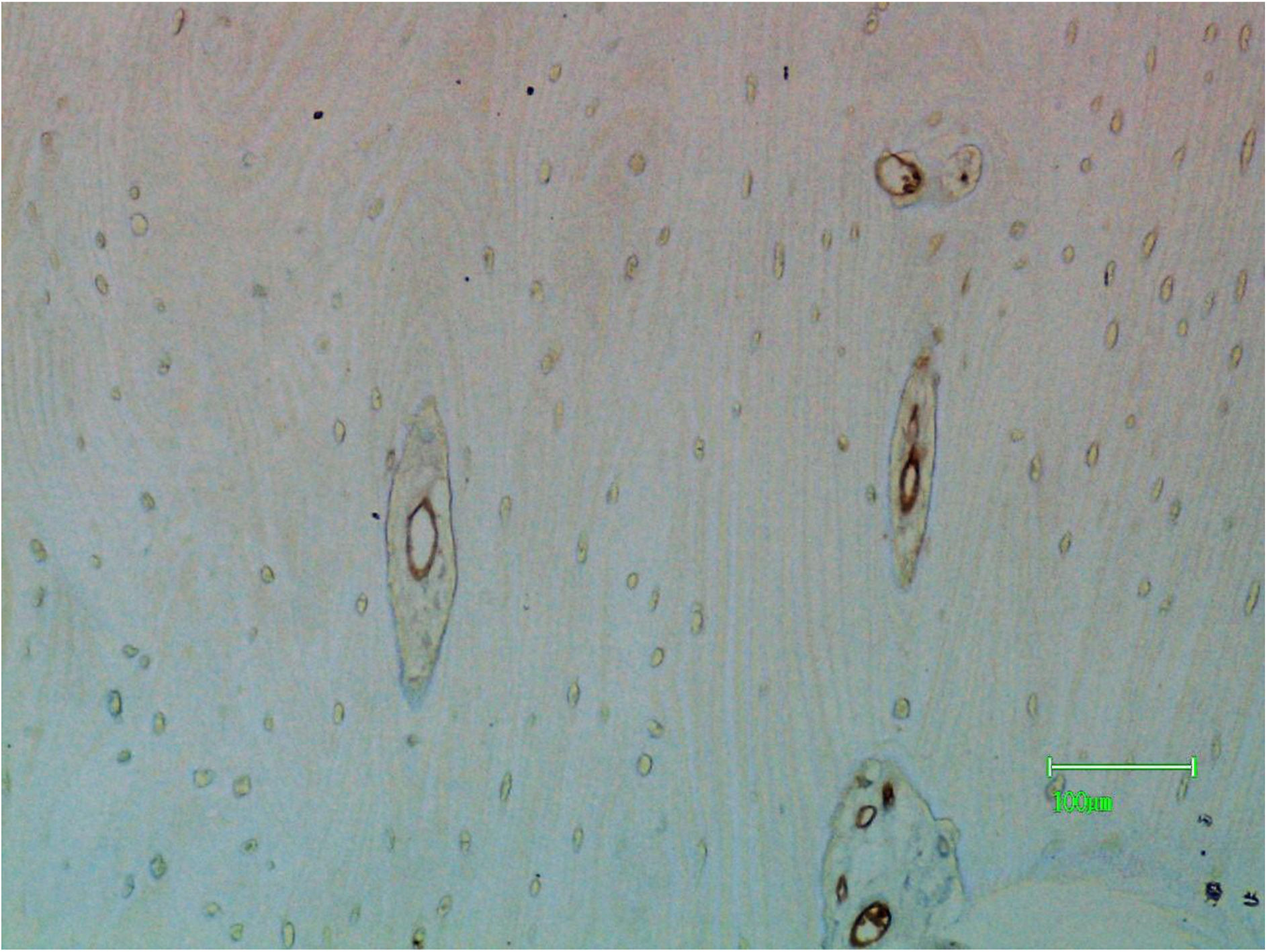
The endothelial cells in cutting channels are positive for PTL-II.

